# CHD4 Interacts With TBX5 to Maintain the Gene Regulatory Network of Postnatal Atrial Cardiomyocytes

**DOI:** 10.1101/2024.12.04.626894

**Authors:** Mason E. Sweat, Wei Shi, Erin M. Keating, Anna Ponek, Jie Li, Qing Ma, Chaehyoung Park, Michael A. Trembley, Yi Wang, Vassilios J. Bezzerides, Frank L. Conlon, William T. Pu

## Abstract

Atrial fibrillation (AF) is the most common sustained arrhythmia, affecting 59 million individuals worldwide. Impairment of atrial cardiomyocyte (aCM) gene regulatory mechanisms predisposes to atrial fibrillation. The transcription factor TBX5 is essential for normal atrial rhythm, and its inactivation causes loss of aCM enhancer accessibility, looping, and transcriptional identity. Here we investigated the mechanisms by which TBX5 regulates chromatin organization. We found that TBX5 recruits CHD4, a chromatin remodeling ATPase, to 33,170 genomic regions (TBX5-enhanced CHD4 sites). As a component of the NuRD complex, CHD4 functions to repress gene transcription. However, combined snRNA-seq and snATAC-seq of CHD4 knockout (KO) and control aCMs revealed that CHD4 has both gene activator and repressor functions. Genes repressed by CHD4 in aCMs included sarcomeric proteins from non-CM cell lineages. Genes activated by CHD4 in aCMs were characterized by TBX5-enhanced CHD4 recruitment, which enhanced chromatin accessibility and promoted the expression of aCM identity genes. This mechanism of TBX5 recruitment of CHD4 was critical for sinus rhythm because *Chd4^AKO^* mice had increased vulnerability to AF from electrical pacing and a fraction had spontaneous AF. Our findings reveal that CHD4 is essential for maintaining aCM gene expression, aCM identity, and atrial rhythm homeostasis.

## Introduction

During each cardiac cycle, an electrical signal emanates from the sinus node, spreads through the atria, and converges on the atrioventricular node, which relays the signal to the ventricles. This electrical signal coordinates the atrial and ventricular contraction, so a single atrial contraction precedes each ventricular contraction. In atrial fibrillation (AF), the most common sustained human arrhythmia, atrial impulse conduction becomes disorganized and forms fractionated, self-sustaining wavelets. This disorganization results in uncoordinated atrial contraction and irregular, often rapid, activation of the ventricles.

Atrial and ventricular cardiomyocytes (aCMs and vCMs) are highly specialized to achieve their distinct roles in the cardiac cycle. These cardiomyocytes have distinct ultrastructures, metabolic activities, and electrophysiological properties, which are maintained and coordinated by the differential expression of thousands of genes. Among these genes are sarcomere components and ion channels that control excitation-contraction coupling and the shape of the CM action potential. Chamber selective gene expression is in turn regulated by a/vCM-specific epigenetic regulatory programs and a/vCM specific chromatin organization^1,2^. Cell type specific differences between aCMs and vCMs are established early in embryonic development and maintained postnatally by distinct gene regulatory mechanisms.

Postnatal aCM identity is controlled by the cardiogenic transcription factor (TF) TBX5, which also regulates development of the atrial and ventricular chambers, conduction system, and interventricular septum^3–5^. In postnatal aCMs, TBX5 inactivation causes loss of chromatin accessibility, the loss of active enhancer mark H3K27ac at aCM enhancers, and the loss of aCM enhancer-promoter loops^6^. Unlike the cardiogenic TF GATA4, which functions as a pioneering TF to directly open chromatin^7^, TBX5 has no known pioneering activity; thus, the ability of TBX5 to maintain enhancer accessibility is likely facilitated by interactions with chromatin remodeling proteins.

In fact, TBX5 interacts genetically and physically with two different chromatin remodelers. The Brg1/Brm-associated factor (BAF) complex facilitates the expression of direct TBX5 target genes *Nppa* and *Bmp10* in mice during development. Compound hemizygous deletions in *Brg1* and *Tbx5* result in severe congenital heart defects, exacerbating the phenotypes of mice with a one-copy of deletion of either gene^8^. TBX5 also physically interacts with chromodomain helicase DNA binding protein 4 (CHD4), the core nucleosome-remodeling enzyme of the Nucleosome Remodelling and Deacetylase (NuRD) complex^9^. The NuRD complex typically represses transcription by deacetylating lysine residues at enhancers. During development, CHD4 deletion in cardiomyocytes resulted in the upregulation of sarcomeric proteins from non-cardiomyocyte muscle lineages^10^, including *Acta1* from skeletal muscle and *Myh11* from smooth muscle tissues. CHD4-dependent inhibition of non-CM lineage genes was facilitated by other TFs, including NKX2-5, GATA4, and TBX5, which recruit CHD4 and NuRD to cis-regulatory elements (CREs) that regulate expression of these genes^11^. This function was shown to be conserved in postnatal cardiomyocytes^12^.

Here, we reveal an unappreciated gene activating function of CHD4 in postnatal aCMs. We show that TBX5 recruitment of CHD4 regulates over 1,000 aCM genes, promoting the expression of ∼500 and repressing ∼600. At the repressed genes, consistent with a prior study, TBX5 recruited CHD4 to prevent genomic accessibility at non-CM lineage genes. In addition to this known function, at the activated genes, CHD4 promoted the accessibility of over 2,000 regions to activate the expression of aCM-selective genes. Nearly half of these regions were bound by TBX5 and exhibited TBX5-enhanced CHD4 binding. The importance of this genetic regulatory mechanism in aCM function was shown by atrial-selective CHD knockout (*CHD4^AKO^*), which caused partially penetrant spontaneous atrial fibrillation and increased vulnerability to pacing-induced atrial fibrillation. Our findings demonstrate that aCM CHD4-mediated gene activation and repression are critical to maintenance of atrial rhythm homeostasis.

## Results

### BRG1 inactivation does not alter postnatal aCM gene regulation

Deletion of TBX5 causes loss of accessibility at enhancer elements that promote aCM-selective gene expression (**Fig. 1a**). Thus, we analyzed the function of the BAF complex component *Brg1,* a TBX5 co-activator^8^. P3 *Brg1^flox/flox^* pups^13^ were injected with an adeno-associated virus serotype 9 (AAV9) encoding *Nppa-Cre* (AAV9:*Nppa-Cre*) or as a control, *Nppa-EGFP* (AAV9:*Nppa-EGFP*) as a control^6,14^. AAV9:*Nppa-Cre* is highly specific to aCMs, and is not active in nodal tissue or vCMs^6,14^. To assess whether inactivating *Brg1* altered TBX5 target gene expression, we performed RTqPCR for previously validated TBX5 target genes on atrial RNA samples from P28 mice, including *Tbx5, Nppa, Nppb, Myl7, Scn5a, Atp2a2,* and *Ryr2* (**Supp. Fig. 1a**)^6,15^. Unexpectedly, *Brg1* inactivation in postnatal aCMs (*Brg1^AKO^*) did not result in the significant downregulation of these TBX5 targets.

**Fig. 1.**
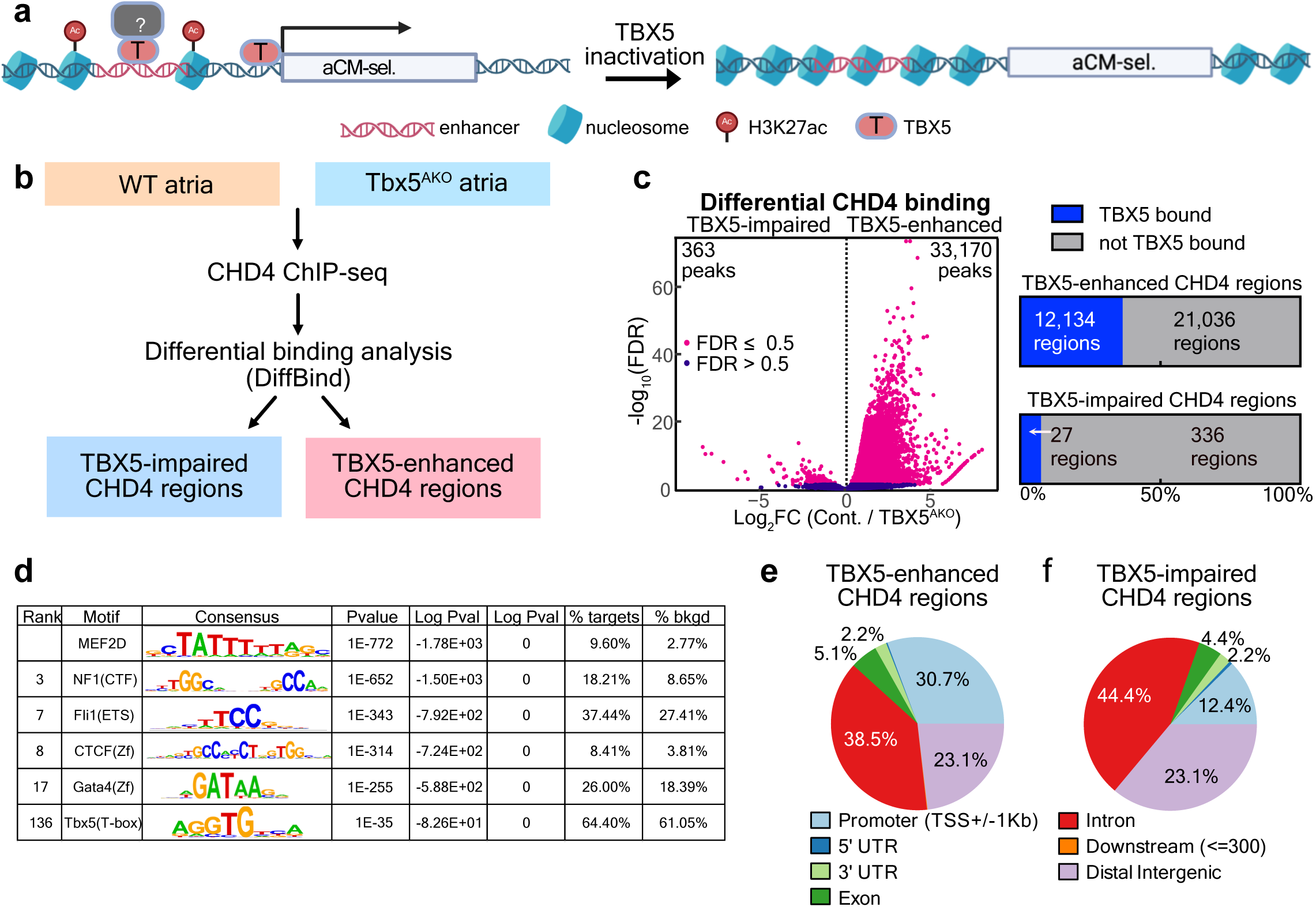
TBX5 recruits CHD4 to genomic loci. **a,** Model of TBX5 gene regulation in aCMs. TBX5 inactivation results in the loss of enhancer accessibility and downregulation of aCM-selective genes. **b,** CHD4 ChIP-seq experimental design. Five animals were used per biological replicate, and two replicates were performed per group. **c,** Diffbind analysis of CHD4 binding in control and Tbx5^AKO^ samples. Significant regions (|Log FC| > 0.5, Log_10_FDR>0.05) are colored red. On the right, intersection of statistically significant regions with TBX5 occupancy data obtained from TBX5 bioChIP-seq in postnatal aCMs (GSE215065). **d,** Motif enrichment analysis of TBX5-enhanced CHD4 regions. **e-f,** Location of TBX5-enhanced and TBX5-impaired CHD4 regions with respect to genome annotations.

To determine whether *Brg1* inactivation altered sinus rhythm in *Brg1^AKO^* mice, we performed surface electrocardiograms (ECG) at P28. In contrast to *Tbx5^AKO^* mice, which develop an AF phenotype by P21^6^, *Brg1* inactivation did not change the cardiac rhythm. The ECG recordings had obvious P waves and regular RR intervals, excluding AF (**Supp.** Fig 1b-c). We concluded that the postnatal aCM gene regulatory network is resistant to *Brg1* inactivation, and we turned our attention to CHD4, another chromatin remodeler that interacts with TBX5^9^.

### TBX5 recruits CHD4 to occupy specific genomic loci

In CMs, CHD4 co-occupies genomic regions with the cardiac TFs GATA4, NKX2-5, and TBX5 near repressed non cardiac muscle cell genes. To determine whether TBX5 actively recruits CHD4 in aCMs, we measured CHD4 occupancy in control aCM nuclei and aCM nuclei from aCM-selective Tbx5 knockout (**Fig. 1b**). Neonatal *Tbx5^flox/flox^* mice were treated as neonates with either AAV9:*Nppa-Cre* (*Tbx5^AKO^*) or AAV9:*Nppa-EGFP* (control) at P3. PCM1 (Pericentriolar Material 1)-positive aCM nuclei were isolated from 3-week-old mice in biological duplicate. We used chromatin immunoprecipitation followed by next generation sequencing (ChIP-seq) to measure CHD4 genomic occupancy. Differential occupancy analysis with Diffbind^16^ identified 33,170 genomic loci that were enriched in control samples compared with *Tbx5^AKO^* (TBX5-enhanced regions, **Fig. 1c, Supp. Table 1**). Overlapping these genomic loci with TBX5 aCM bio-ChIP-seq data^17^ demonstrated that 12,134 (36.6%) of the 33,170 regions were directly bound by TBX5 in postnatal aCMs. In contrast to this large number of TBX5-enhanced CHD4 regions, far fewer CHD4 regions had increased CHD4 binding following TBX5 KO (363 TBX5-impaired CHD4 regions). We found that 60,810 (64.5%) CHD4 binding sites did not show significant enrichment in either control or KO samples.

To identify regulators of the TBX5-enhanced CHD4 binding regions, we performed motif enrichment analysis. The top-most enriched motif corresponded to MEF2, a well-characterized cardiomyocyte transcription factor^18^ (**Fig. 1d**). TBX5-enhanced binding regions were also enriched in the motifs of cardiac TFs GATA4 and TBX5, and for CTCF, which was previously associated with CHD4 in CHD4-IP/Mass Spectrometry (MS) experiments in cardiomyocytes^19^, specifically in regions associated with GATA4, NKX2-5, and TBX5^11^. MEF2, GATA4, NKX2-5 are established TBX5 interacting proteins^20–22^.

The genomic distribution of the TBX5-enhanced CHD4 binding regions revealed that a disproportionate number of regions were within gene promoters (30.7%, +/− 1kb of the TSS). Other common locations included introns (38.5%) and distal intergenic regions (23.14%) (**Fig. 1e**). Proportions of binding locations of TBX5-depleted regions were similar in frequency, with the exception of promoter regions (12.4%) (**Fig. 1f**).

Together these data indicate that TBX5 recruits CHD4 to genomic binding sites, particularly promoters, co-occupied by TBX5 and other cardiac TFs.

### CHD4 inactivation results in atrial remodeling and immune cell infiltration

To identify effects of CHD4 inactivation in postnatal aCMs, we injected AAV9:*Nppa-Cre* and AAV9:*Nppa-EGFP* into P3 *Chd4^flox/flox^* pups (**Fig. 2a**). Echocardiography parameters were not significantly different between mice treated with control or AAV9:*Nppa-Cre* at 1 to 3 months (**Fig. 2b**). To confirm CHD4 inactivation, heart sections from 2-3 month old mice were stained for CHD4 and either fibronectin type II and SPRY domain containing protein 2 (FSD2) or sarcomeric alpha actinin (SAA), markers of cardiomyocytes^23^(**Fig. 2c**). Control hearts had robust staining for CHD4 in aCM and non-aCM nuclei; however, *Chd4^AKO^* hearts had clear loss of CHD4 in aCMs. Because the activity of the *Nppa* promoter is restricted to aCMs, non-CM cells in *Chd4^AKO^* atria retained CHD4 staining within their nuclei (white arrowheads, Fig. 2c). No difference in CHD4 staining was observed in ventricular sections (**Fig. 2d**).

**Fig. 2.**
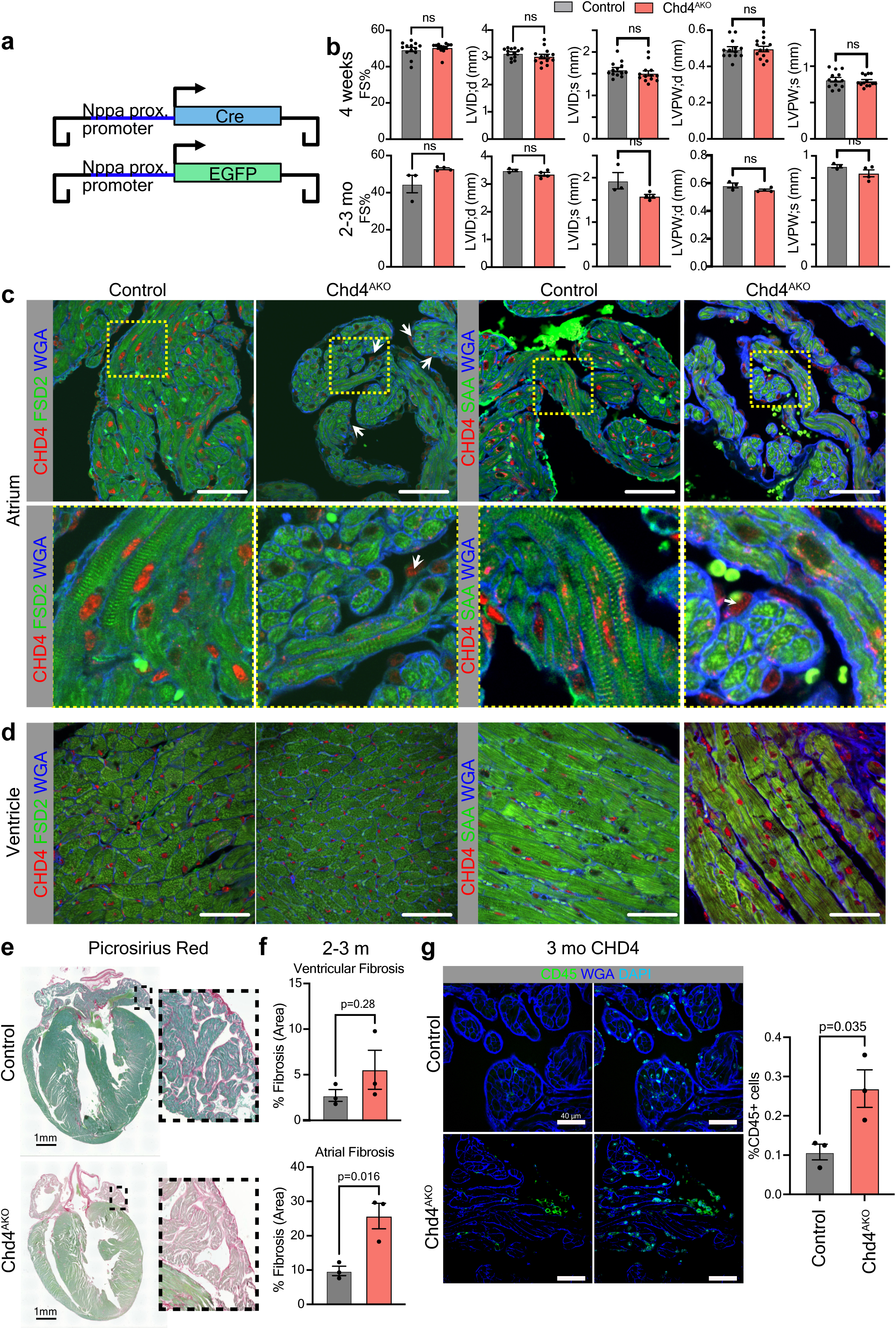
Inactivation of CHD4 in aCMs results in atrial remodeling. **a,** Schematic of the AAV9:Nppa-EGFP and AAV9:Nppa-Cre AAV constructs. Chd4^flox/flox^ animals that received AAV9:Nppa-EGFP and AAV9:Nppa-Cre were designated control and Chd4^AKO^, respectively. **b,** Echocardiographic assessment of left ventricular size and function. Studies were performed 4 weeks and 2-3 months after AAV injection. 4 weeks, n=13 per group. 2-3MO, n=3 control and 4 Chd4^AKO^. Two tailed t-test, ns, P>0.05. **c-d,** Confocal images of immunostained atrial and ventricular sections. White arrows in the Chd4^AKO^ images denote cells that retained CHD4 expression. Yellow boxed areas are enlarged in the third column of panels. **e,** Picrosirius red staining of control and Chd4^AKO^ heart sections. Boxed areas are enlarged to the right. **f,** Quantitation of fibrotic area. n=3 mice per group. Unpaired t-test. **g,** Confocal images of immunostained atrial sections from 3-month-old mice. Right, quantification of the percentage of CD45+ cells per section compared to the total number of cells. n=3 mice per group. Unpaired t-test.

Fibrotic remodeling is a hallmark of abnormal atrial function. To determine whether CHD4 inactivation resulted in atrial fibrosis, we performed picrosirius red/fast green staining in 2-3 month old sections of control and *Chd4^AKO^* hearts (**Fig. 2e-f**). *Chd4^AKO^* mice had strikingly elevated levels of atrial fibrosis, whereas there was no difference in ventricular fibrosis.

In humans and mouse models of AF, Hulsmans *et al.* noted increased immune cell recruitment to atria^24^. To determine whether inactivation of CHD4 resulted in increased numbers of immune cells, we performed staining for the pan-immune cell marker CD45 (**Fig. 2g**). *Chd4^AKO^*atria had greater numbers of CD45+ cells compared with controls.

### CHD4 and TBX5 coordinately regulate atrial gene expression

The recruitment of CHD4 by TBX5 to genomic loci suggested that TBX5 and CHD4 coordinately regulate gene expression in postnatal aCMs, similar to what occurs for vCMs during development^10,11^. To test this hypothesis, we performed bulk RNA-seq of the right and left atria of P21 *Chd4^flox/flox^* mice treated with either AAV9:*Nppa-Cre* or AAV9:*Nppa-EGFP* at P3 (n=4/group; **Fig. 3a**). Principal component analysis demonstrated a clear separation of the control samples from *Chd4^AKO^*along PC1, which included 70% of the variance for the assay (**Fig. 3b**). DEseq2 analysis identified 2,748 differentially expressed genes ( | Log_2_FC | > 0.5 and P_adj_ <0.05), including 1,077 with greater expression in controls and 1,671 with greater expression in *Chd4^AKO^* (**Supp. Table 2, Fig. 3c**).

**Fig. 3.**
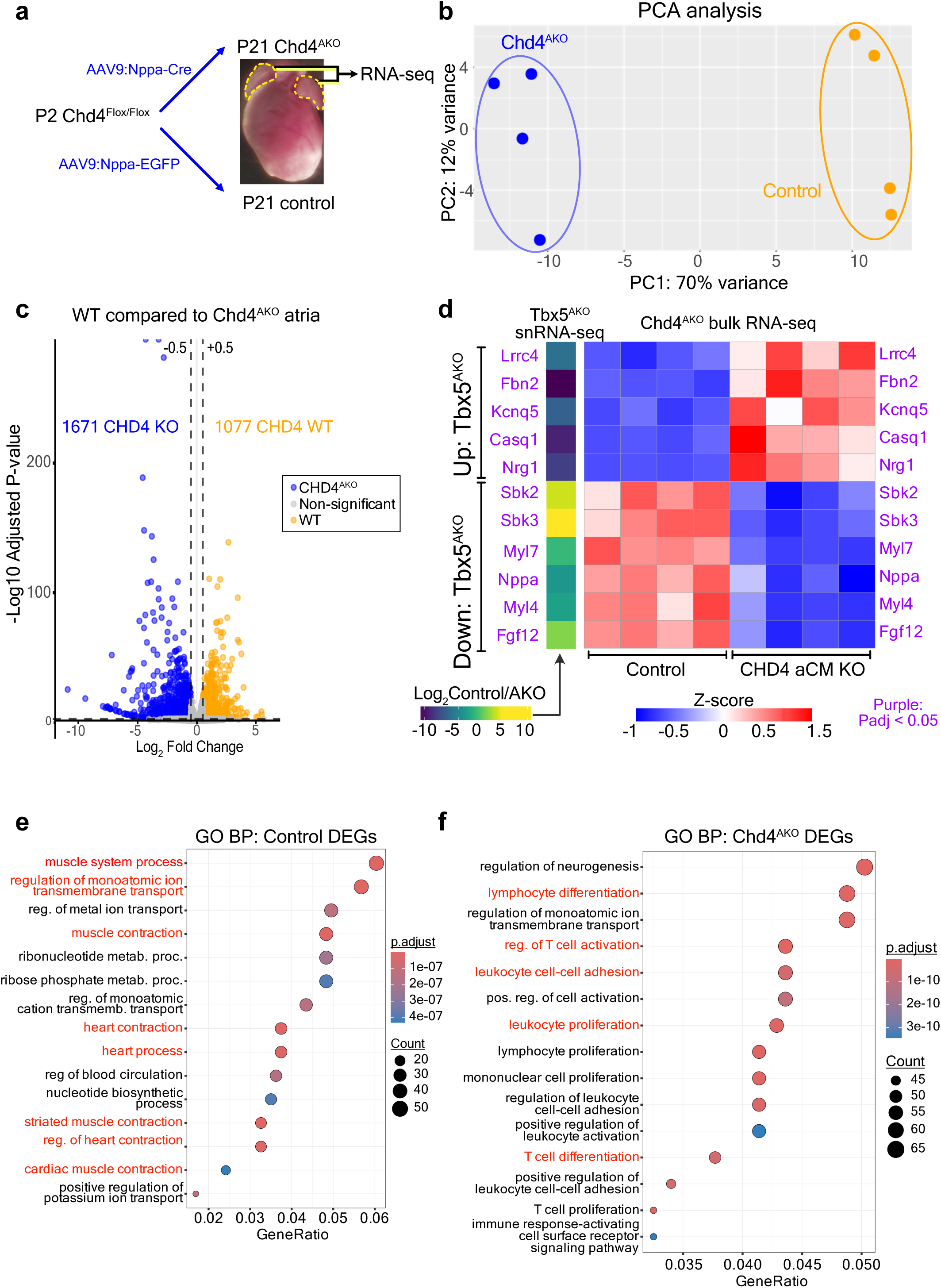
Bulk RNA-sequencing of P21 control and Chd4^AKO^ atria. **a,** Experimental design. P21 Right and left atria from four control mice and four Chd4^AKO^ mice were used in the experiment. **b,** Principle component analysis of WT and Chd4^AKO^ RNA-seq samples. **c,** Differentially expressed genes between WT and Chd4^AKO^ atria. Significantly differentially expressed genes (DEGs) had |Log_2_FC|>0.5 and Adjusted P-value<0.05. **d,** Heatmap of aCM-selective TBX5 target genes and genes upregulated in TBX5 KO aCMs in snRNA-seq data from WT and Tbx5^AKO^ aCMs compared to WT and Chd4^AKO^ atria. Gene symbols in purple were significantly differentially expression in control vs KO in both Tbx5 and Chd4 knockout experiments. **e,f,** GO enrichment terms enriched for control or Chd4^AKO^ DEGs. The X axis shows the ratio of the number of genes in the intersection of DEGS and the gene set to the total genes in the gene set. Datapoints are colored by the adjusted P value, and datapoint size corresponds to the total number of genes for each term. GO terms highlighted in red are of particular biological interest.

To determine whether CHD4 regulated TBX5 target genes, we examined a curated list of aCM-selective genes that are downregulated in *Tbx5^AKO^* aCMs and regulated by TBX5-dependent chromatin looping (*Sbk2, Sbk3, Myl7, Nppa, Myl4* and *Fgf12*)^6^, and a list of genes that are upregulated in *Tbx5^AKO^* aCMs (*Lrrc4, Fbn2, Kcnq5, Casq1* and *Nrg1*). Expression changes in this gene set were examined in our previously reported *Tbx5^AKO^* snRNA-seq dataset^6^ and the *Chd4^AKO^* bulk RNA dataset (**Fig. 3d**). This analysis revealed that for these pre-selected targets, gene expression changes between WT and *Chd4^AKO^* atria were largely comparable to differences between WT to *Tbx5^AKO^*aCMs. GO term analysis revealed that genes more highly expressed in control CHD4 samples were correlated with terms related to muscle function (**Fig. 3e**). Genes upregulated in *Chd4^AKO^* samples were more related to lymphocyte activation and atrial remodeling (**Fig. 3f**), a result consistent with the atrial remodeling, fibrosis, and immune cell infiltration that we observed in *Chd4^AKO^* atria (**Fig. 2e-g**).

To extend the analysis to all of the differentially expressed genes (DEGs, P_adj._<0.05) detected in the *Chd4^AKO^* bulk RNA-seq and the TBX5 snRNA-seq assays, we compared the change in each gene’s expression caused by aCM-selective ablation of *Chd4* or *Tbx5* (**Supp.** Fig 2a). Most DEGs common to both experiments were altered in the same direction by TBX5 or CHD4 inactivation (purple points in quadrants I and III). In each dataset, genes with aCM-selective expression were more highly expressed in the WT compared with the AKO group (**Supp.** Fig 2b), consistent with CHD4 and TBX5 promoting their expression. In total, CHD4 and TBX5 coordinately promoted the expression of 513 genes and repressed the expression of 633 genes (**Supp. Fig. 2c,d**). The degree of overlap between DEGs downstream of CHD4 or TBX5 was highly significant (Hypergeomtric test: P<2.5E-34). GO term enrichment for DEGs common to both experiments revealed that genes activated by both factors facilitate metabolism and nucleotide biosynthesis, whereas genes repressed by both CHD4 and TBX5 were related to extracellular matrix remodeling and TGFβ signaling (**Supp. Fig. 2e**).

### CHD4 inactivation increases atrial fibrillation vulnerability

Because CHD4 and TBX5 cooperatively regulate target genes, and that the inactivation of TBX5 in aCMs causes AF^6^, we performed surface ECG measurements on *Chd4^AKO^* and control mice to examine atrial rhythm. Rhythm regularity was assessed by the standard deviation of the RR interval (SDRR; **Fig. 4a**). At ages 1 and 2-3 months, there was no statistically significant elevation in SDRR, suggesting that mice remained in a regular rhythm. At 4-5 months old, there was a tendency toward elevated SDRR (p=0.051); one 4-5 week-old and three 5 month-old mice had markedly elevated SDRR and no P waves, hallmarks of AF. Therefore, 4 out of 15 *CHD4^AKO^* mice examined by surface ECG across different timepoints had spontaneous, irregular rhythms consistent with AF, compared with zero of 12 in the control group (**Fig. 4b**).

**Fig. 4.**
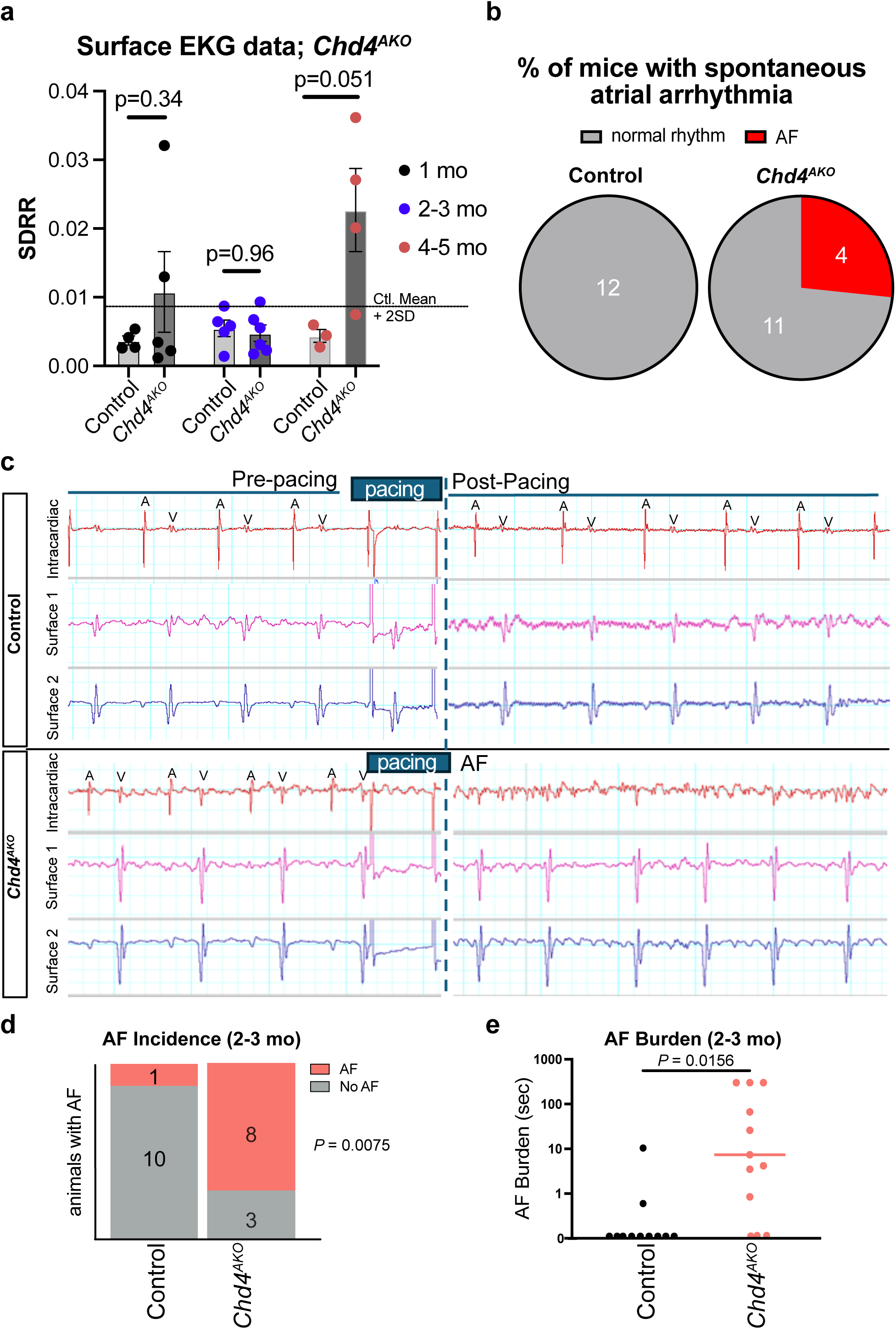
*Chd4* inactivation in aCMs increases AF vulnerability. **a**, Irregularity of cardiac rhythm was assessed from surface EKGs using the standard deviation of the R-R interval (SDRR). While SDRR was not significantly elevated at any timepoint, four mice (one at 4-5WO and 3 at 4-5MO) had elevated SDRR. Surface ECGs of these mice showed a loss of P waves, consistent with AF. Unpaired *t*-test. The dotted line represents the upper limit of normal for control mice (mean + 2SD). **b**, The total fraction of mice with spontaneous atrial arrhythmias. **c**, Representative data from a programmed atrial stimulation experiments. The atrial rhythm was recorded by an intracardiac electrode, and the surface ECG was captured by limb leads. Following the pacing protocol, the atrial recording was evaluated for AF. A, atrial signal. V, ventricular signal. **d**, AF incidence following programmed atrial simulation of 2-3 mo mice. 1/11 control and 8/11 *Cdh4^AKO^* had AF for greater than 3 seconds following pacing, while 8 of 11 *Cdh4^AKO^* mice exhibited atrial arrhythmia. Fisher’s exact test. **e,** AF burden from the 2-3MO cohort of paced mice. AF burden was significantly higher in *Cdh4^AKO^* mice. Wilcoxon test.

Intracardiac pacing of the right atrium is a common approach to examine AF vulnerability in mice. With a pacing protocol used for TBX5 heterozygous mice^25^, we tested the AF vulnerability of CHD4^AKO^ mice at 2-3 months old (**Fig. 4c**). A greater proportion of *Chd4^AKO^* could be induced to develop sustained AF (lasting greater than 3 seconds, Fisher’s exact test *P*=0.0075) compared to control (**Fig. 4d**). The duration of induced AF was also increased in *Chd4^AKO^* mice, including three mice that did not recover normal rhythm by the termination of the study five minutes after programmed stimulation (**Fig. 4e**).

### Multiomic analysis of control and Chd4^AKO^ mice atria

Given the canonical function of CHD4 in chromatin remodeling, we performed concurrent single nucleus RNA-seq and ATAC-seq (multiomics) on control and *Chd4^AKO^* atria. In total, four *Chd4^AKO^* samples and two control samples were processed, with each sample consisting of a right and left atrium from two mice. Two *Chd4^AKO^* samples were obtained at 3.5 months from mice that exhibited AF upon electrical pacing, and two *Chd4^AKO^*samples were obtained at 5 months from mice, three mice of which exhibited spontaneous atrial arrhythmia (**Supp. Table 3**). Wild-type samples were obtained from 5 month old controls. Each sample consisted of a male and female mouse. Following doublet removal and rigorous quality control for both the RNA and ATAC data (see Methods), we obtained a total of 27,263 high quality nuclei (**Supp. Fig. 4a**). These nuclei from the six samples were merged into a single Seurat object (Methods), and the snRNA, snATAC data, and weighted nearest neighbors (WNN) combination of both types of data were used to create Uniform Manifold and Approximation Projection (UMAP, **Supp. Fig. 3a**) embeddings. **Supp. Fig. 4a and b** show final sample metrics for the RNA and ATAC datasets. Louvain clustering was performed on the WNN integrated data to identify the major cell types. Metadata from the originating sample and sample genotype were used to split the WNN UMAP, revealing high concordance between samples of each genotype and an overall high concordance with minimal batch effect among all the datasets (**Supp. Fig. 4c,d**).

Initial cell state determinations, made using SC-type^26^, were refined using marker genes for different atrial cell types (identity genes, **Supp. Fig. 3c**). The number of nuclei and the normalized contribution from *Chd4^AKO^* and WT samples were determined (**Supp. Fig. 3b,c**). This annotation process enabled the removal of a small cell cluster with high expression of epicardial and immune cell markers, likely residual doublets. Annotation also revealed cell states that were highly enriched in either *Chd4^AKO^* or control samples, including aCM_1, the largest aCM cluster containing predominantly control nuclei. Smaller aCM clusters aCM_3, aCM_4, and aCM_5 mostly contained control nuclei. In contrast, the second largest aCM cluster, aCM_2, was nearly exclusively derived from *Chd4^AKO^*samples. There was also genotype-specific enrichment for endocardial (Endocardium_3, control; Endocardium_2, *Chd4^AKO^*) and macrophage clusters (Macrophage_3, control, Macrophage_2, *Chd4^AKO^*).

### CHD4 controls the aCM identity gene program

To identify genes differentially expressed between control and *Chd4^AKO^* aCMs, we compared aCM_1 and aCM_2, the largest aCM clusters that predominantly originated, respectively, from control and *Chd4^AKO^* samples (**Fig. 5a,b**). We identified 2,256 DEGs (|Log_2_FC| > 0.5 and P_adj_<0.05), of which 431 and 1825 were higher in aCM_1 and aCM_2, respectively (**Supp. Table 4**). DEGs from this snRNAseq comparison agreed with those from the bulk RNA-seq dataset of control and *Chd4^AKO^* atria (Pearson r=0.835; P=1.9E-302. **Supp. Fig. 5a**). GO terms for genes upregulated in control and *CHD4^AKO^* aCMs were similar to the GO terms identified in the bulk RNA-seq experiment (**Fig. 5c**); terms enriched for *Chd4^AKO^*DEGs are related to chamber remodeling, while terms enriched for control DEGs suggest important roles for muscle function, cell junction assembly and cardiac rhythm.

**Fig. 5.**
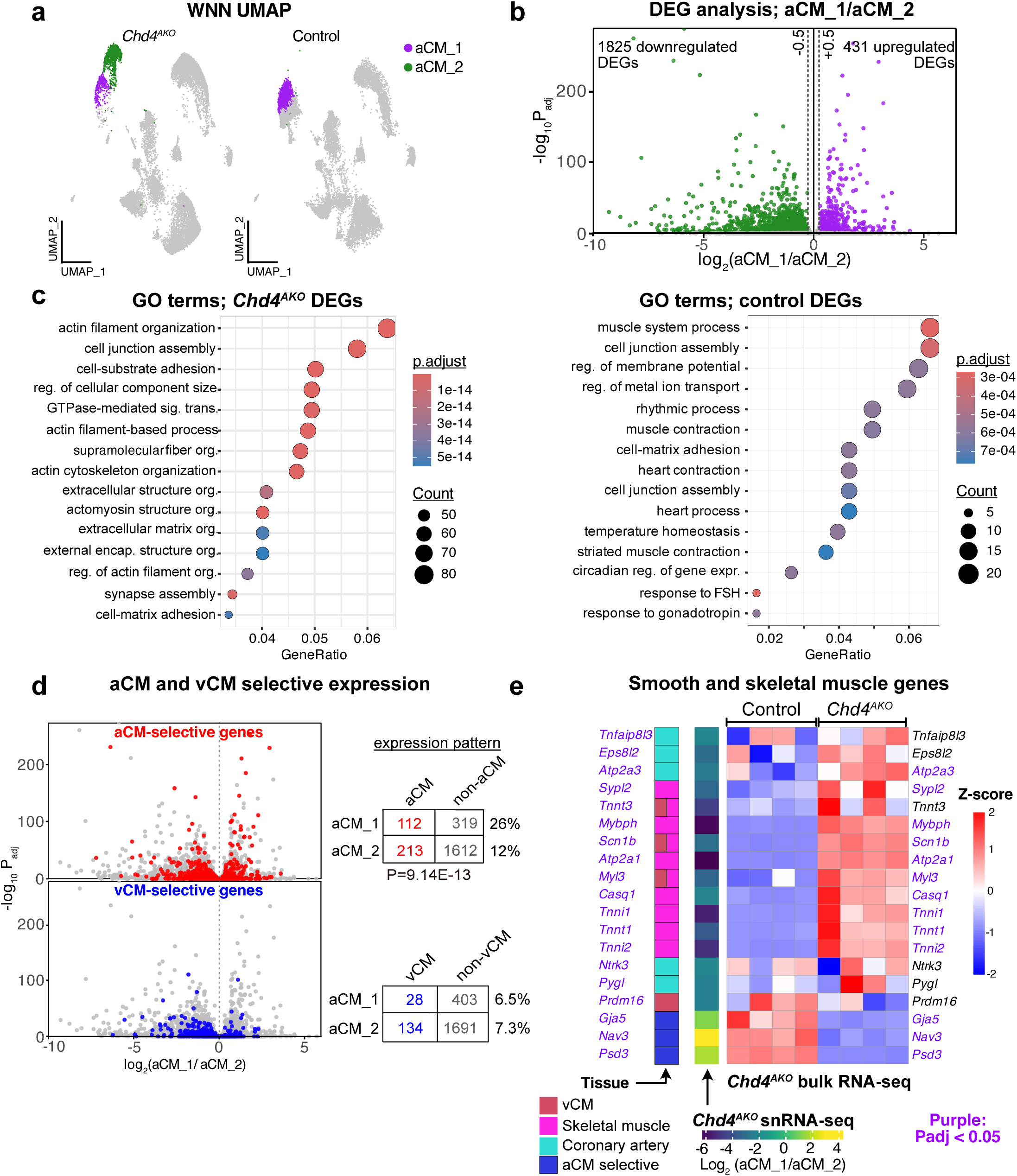
Combined snRNA-seq and snATAC-seq identifies distinct WT and *CHD4^AKO^* cell states aCMs. **a**, Concurrent snRNA-seq and snATAC-seq comparison of *Chd4^AKO^* and control atria. Weighted nearest neighbor (WNN) UMAP integration of both data types is split by sample type. aCM_1 and aCM_2 predominantly originated from *Chd4^AKO^* and control atria, respectively. **b**, Pseudobulk RNA-seq analysis of genes differentially expressed (|Log2FC|>0.5 and Adjusted P-value <0.05) between aCM_1 (control) and aCM_2 (*Chd4^AKO^*). **c**. Gene ontology germs enriched for DEGs more highly expressed in *Chd4^AKO^* or control. **d**, Altered expression of aCM and vCM selective genes in *Chd4^AKO^* or control aCMs. Red and blue dots indicate aCM-selective and vCM-selective DEGs, respectively, as defined in Cao et al. 2023 (GSE215065). Contingency tables for Fisher test are shown to the right. **e**, Expression of smooth muscle and skeletal muscle genes in *Chd4^AKO^* and control aCMs. The left-most column indicates normal tissue-selective expression, based on GTEX (**Supp. Fig. 5**). The next column indicates fold-change in gene expression from control vs. *Chd4^AKO^* aCMs. The main heatmap indicates row-scaled expression in bulk RNA-seq from *Chd4^AKO^* versus control atria. Genes colored purple were significantly differentially expressed in the snRNA-seq or bulk RNA-seq experiments, respectively.

To determine whether CHD4 regulated the atrial identity program, aCM-selective and vCM-selective genes were overlaid with DEGs from the CHD4^AKO^ vs control aCM (aCM_2 vs aCM_1) comparison (**Fig. 5d**). Atrial CM-selective genes were overrepresented among genes activated vs repressed downstream of CHD4 (26% (112/413) vs. 12% (213/1825); Fisher’s exact test P=9.1E-13). In contrast, we did not observe significant enrichment of vCM-selective genes among genes activated or repressed downstream of CHD4.

In fetal and postnatal cardiomyocytes, CHD4 interacts with cardiac transcription factors including TBX5 to repress the expression of skeletal and smooth muscle lineage genes^10,12^. To determine whether this function is conserved in aCMs, we examined top 200 DEGs ranked by significance in control and *Chd4^AKO^* aCMs for their expression levels in the left ventricle, atrial appendage, skeletal muscle and coronary arteries from GTEX, a human RNA-seq database^27^ (**Fig. 5e** and **Supp.** Fig 5b). The top-ranked DEGs activated downstream of CHD4 tended to be highly expressed in atrial appendage and ventricle, with few showing enrichment in coronary artery or skeletal muscle. In contrast, top-ranked DEGs repressed downstream of CHD4 were enriched in skeletal muscle genes (e.g., *Myl3, Tnnt3, Tnnt1, Tnni1, Atp2a1, Acta1, Casq1, Ypl2, Scn1b*) or in coronary artery genes (e.g., *Prdm16, Eps8l2, Pygl, Atp2a3*). Heatmaps generated from the CHD4 snRNA-seq and bulk RNA-seq datasets (**Fig. 5e**) confirmed the downregulation of aCM-selective genes and the upregulation skeletal and coronary artery tissue genes. These data revealed that CHD4 is critical for preventing the misexpression of genes from other muscle lineages in postnatal aCMs.

### A dual function for CHD4 in enhancer accessibility maintenance and repression

Having identified and validated by multiple approaches the identity of *Chd4^AKO^* and control aCMs in the multiome dataset, we examined differences in chromatin accessibility between *Chd4^AKO^*and control aCMs (**Fig. 6a**). There were 13,935 differentially accessible regions (DARs) between control and *Chd4^AKO^* aCMs (aCM_1 vs. aCM_2), with 2,471 and 11,464 DARs with significantly greater accessibility in control (control DARs) and *Chd4^AKO^* (*Chd4^AKO^* DARs), respectively (**Supp. Table 5**). Most control DARs (83.6%; 2,061 regions) overlapped CHD4 regions, suggesting that CHD4 directly participates in maintaining the accessibility of most of these regions. In contrast, only 41.5% (4,760 regions) of *Chd4^AKO^* DARs were occupied by CHD4, suggesting that CHD4 governs the accessibility of these regions by a mix of direct and indirect mechanisms. Nevertheless, twice as many DARs directly occupied by CHD4 gained, instead of lost, accessibility with CHD4 depletion, consistent with the canonical NuRD complex function of CHD4 to deacetylate and repress enhancers^28^. To determine whether the control and *Chd4^AKO^* DARs regulate aCM or vCM-selective genes, the nearest gene to each DAR was assigned as the putative regulatory target of the DAR (**Fig. 6b**). This analysis revealed that 15.7% of control DARs neighbor aCM-selective genes, while only 4.2% neighbor vCM-selective genes. 11.1% of *Chd4^AKO^* DARs neighbored aCM-selective genes, and 6.2% neighbored vCM-selective genes.

**Fig. 6.**
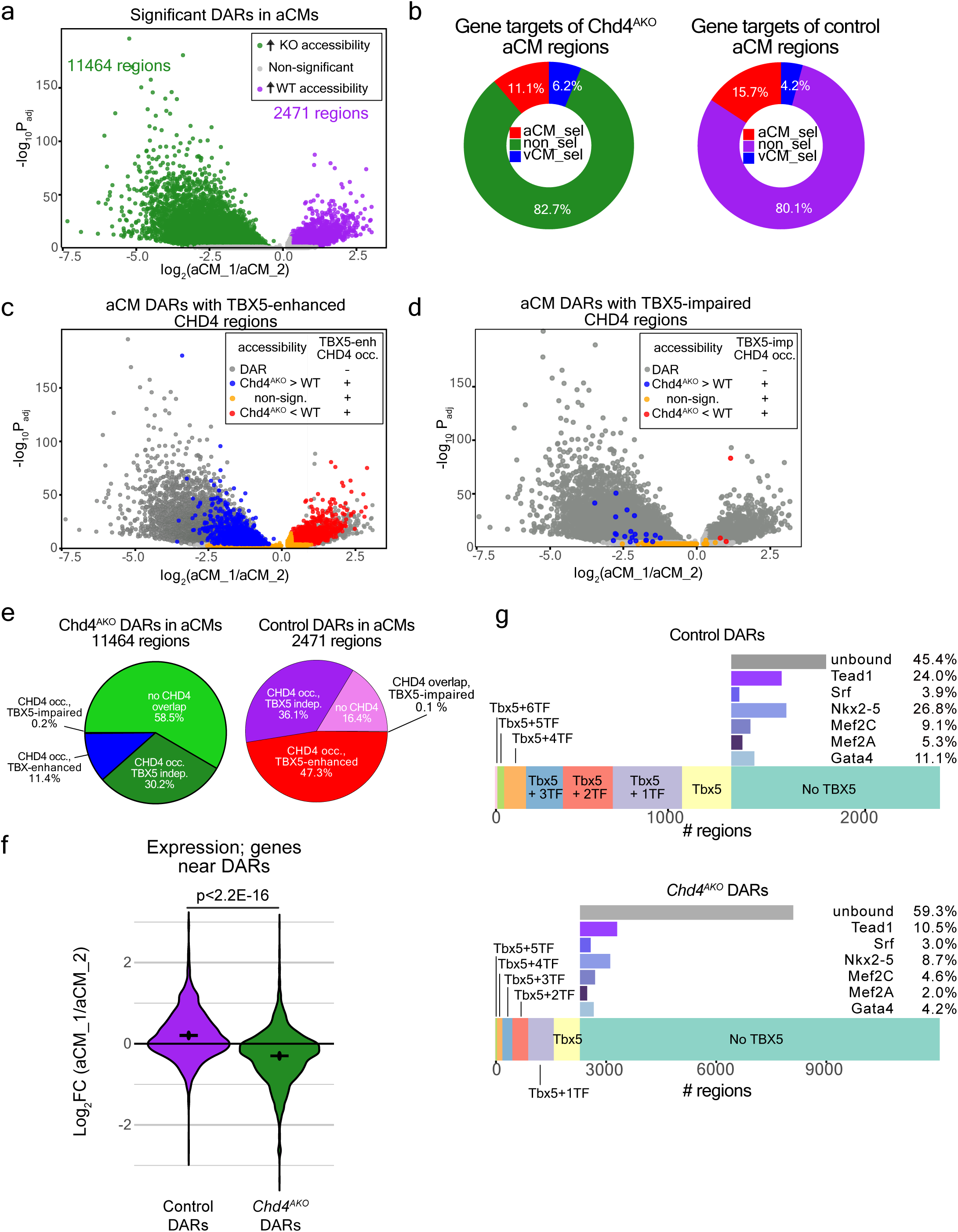
Transcriptional activation by CHD4. **a,** Differentially accessible regions (DARs) in control vs. Chd4^AKO^ aCMs. DARs were defined by | Log_2_ FC|>0.25 and P_adj_<0.05. **b,** The proportion of control and Chd4^AKO^ DARs neighboring aCM or vCM-selective genes. **c-d,** DARs were colored by their overlap with TBX5-enhanced (c) and TBX5-impaired (d) CHD4 regions. **e,** The proportion of control and Chd4^AKO^ DARs overlapping with CHD regions. CHD4 regions were divided into TBX5-enhanced, TBX5-impaired, and TBX5-independent subsets. **f,** Difference in aCM transcript levels of genes neighboring control or Chd4^AKO^ DARs. Included genes were detected in ≥5% of aCM_1 or aCM_2 nuclei. The lines on the plot represent mean+/− SEM. Wilcoxon rank-sum test. **g,** TF binding patterns in wild-type aCMs at control and Chd4^AKO^ DARs. The left hand part of the bar represents regions that are bound by TBX5, stratified by the number of additional binding factors. The right hand portion of the bar represents regions not bound by TBX5, showing the proportion of regions bound by each indicated TF, or unbound by any of the profiled TFs.

In aCMs, TBX5 recruited CHD4 to 33,170 genomic loci (**Fig. 1c**). These Tbx5-enhanced CHD4 regions overlapped 1,168 control DARs and 1,302 *Chd4^AKO^* DARs (**Fig. 6c**). The same analysis using TBX5-impaired CHD4 regions revealed only 4 and 19 regions with greater accessibility in control or *Chd4^AKO^*, respectively (**Fig. 6d**). Strikingly, 47.3% (1,168/2,471) of control DARs used TBX5 to facilitate CHD4 recruitment, compared with only 11.4% (1,302/14,464) of *Chd4^AKO^* DARs (**Fig. 6e**). We validated these results by analyzing reported H3K27ac ChIP-seq data from WT and *Tbx5^AKO^*aCMs^6^. Control DARs exhibited modest H3K27ac signal in WT aCMs, in agreement with their enhancer activity (**Suppl. Fig. 7a**). Control DAR H3K27ac and chromatin accessibility were reduced by *Tbx5* inactivation, consistent with Tbx5-dependent activation of these enhancers. The comparable analysis of *Chd4^AKO^* DARs was complicated by the low accessibility and enhancer activity of these regions in WT aCMs, i.e., there was little detectable H3K27ac signal (**Suppl. Fig. 7a**). Also, 88.5% of these regions were not TBX5-dependent, and their chromatin accessibility was increased only in *Chd4^AKO^* and not *Tbx5^AKO^* aCMs.

To determine how differential accessibility of these regions affected gene expression, we measured the transcript levels of genes in proximity to control regions and KO regions in control and *Chd4^AKO^* aCMs (**Fig. 6f**). As a group, genes near control regions were expressed at greater levels in control aCMs, and genes near KO regions were expressed at greater levels in *Chd4^AKO^* aCMs (Wilcoxon rank-sum test, *P*<2.2E-16).

We created a diagram of the control and *Chd4^AKO^* TBX5-enhanced CHD4 bound DARs (**Supp. Fig. 6a,b**), detailing if the DAR neighbored an aCM-selective, vCM-selective, or non-selective gene, and if the region was associated with a DEG regulated by CHD4 in the snRNA-seq experiment. Of the 1,168 TBX5-enh., CHD4 bound control DARs, 206 neighbored an aCM-selective gene, and of these 206 regions, 97 (47%) were associated with an aCM-selective gene with increased expression in control aCMs. A small number of the 206 regions neighbored aCM-selective genes (13, 6%) repressed by CHD4, and the rest (98 regions, 48%) neighbored aCM-selective genes insensitive to CHD4 inactivation. Few of the 1,168 TBX5-enh., CHD4 bound control DARs were associated with vCM-selective genes (40). The majority of the 1,168 control TBX5-enhanced CHD4 bound DARs (922) neighbored genes that did not show aCM or vCM selective expression, but only 127 (14%) of the 922 regions neighbored genes enhanced by CHD4, and 70 (8%) neighbored genes repressed by CHD4. Interestingly, non-selective genes enhanced by TBX5-enhanced CHD4 bound DARs included genes with important functions in cardiomyocytes, including *Ryr2, Rbm20,* and *Cilp1/2,* amongst others.

The same analysis of the 1,302 TBX5-enhanced, CHD4 bound *Chd4^AKO^* DARs revealed a greater number of regions (79 regions) associated with vCM-selective genes; 36 regions neighbored vCM-selective genes repressed by CHD4. Of the 1,302 DARs which were associated with aCM-selective genes (162), 59 were associated with aCM-selective genes repressed by CHD4. The repressed aCM-selective genes included genes such as *Atp2a1,* and *Myl1,* which are expressed at greater levels in aCMs than vCMs but are more often associated with skeletal muscle, and *Tgfb2*, associated with chamber remodeling.

Next, we examined how aCM-selective, vCM-selective, and non-selective genes associated with the three major types of DARs identified by our analysis (TBX5-enhanced CHD4 bound DARs; CHD4 bound DARs; and DARs which did not bind to either TBX5 or CHD4, **Supp. Fig. 6c**) changed in control and *Chd4^AKO^* aCMs. CHD4 occupied, TBX5-enh. DARs had a greater activating role than CHD4 bound DARs and DARs lacking CHD4 and TBX5 for non-selective and aCM-selective genes. These trends were evident, but not significant, in vCM-selective and skeletal muscle selective gene sets. Notably, a number of skeletal muscle selective genes that are highly upregulated in *Chd4*^AKO^ aCMs are controlled by DARs lacking TBX5 and CHD4.

Together, these data reveal dual repressive and activating functions of CHD4 in aCMs. In agreement with prior studies^10^, TBX5 recruited CHD4 to a set of genomic loci to repress gene expression, and CHD4 recruited by TBX5 also activated a subset of aCM-selective genes.

### CHD4 co-occupancy with other cardiac transcription factors

TBX5 is only one of several TFs that have important functions in aCM gene regulation. CHD4 interacts with some of these other TFs, including GATA4 and NKX2-5^11^. We used previously generated BioChIP-seq data^2^ to examine the occupancy of TBX5, GATA4, MEF2A, MEF2C, NKX2-5, SRF and TEAD1 at *Chd4^AKO^* and control DARs in WT aCMs using previously generated BioChIP-seq data^2^ (**Fig. 6g**). TBX5 was the most prominently bound TF at both control and *Chd4^AKO^* DARs; TBX5 bound 53% of control DARs. NKX2-5 and GATA4, known CHD4-interacting TFs, co-occupied 26.8% and 11.1% of control DARs, respectively. TEAD1 co-bound 24.0% of control DARs. These data suggested that TBX5 is the primary factor that recruits CHD4 as a potential activator, with NKX2-5, TEAD1, and GATA4 being additional candidate TFs. A smaller percentage of *Chd4^AKO^* DARs were bound by TBX5 (19%) or other cardiac TFs, suggesting that CHD4 recruitment to repressed loci predominantly occurs through mechanisms independent of binding to the major cardiac TFs.

To identify TFs that also participate in CHD4 recruitment, we performed motif enrichment analysis on control and *Chd4^AKO^* DARs that were bound by CHD4 in WT aCMs (**Supp. Fig. 6b**; full list in **Supp. Table 6**). Many motifs (91) were enriched in both control and *Chd4^AKO^* DARs, including motifs for TBX5, MEF2A/C, and GATA2/4/6. We focused on motifs uniquely enriched in either control or *Chd4^AKO^* DARs. We found 30 motifs enriched in control DARs and 147 motifs in *Chd4^AKO^* DARs. The top uniquely enriched motif in control DARs belonged to GR, the glucocorticoid receptor, which is implicated in cardiac fibrosis by interactions with the mineralocorticoid receptor (NR3C2), another control-enriched motif^29^. Top uniquely enriched motifs in *Chd4^AKO^*DARs belonged to Fos, a well-characterized protooncogene^30^, and ETV2, a master regulator of vascular development^31^.

### CHD4 regulates atrial fibrillation associated genes

Our data indicated that CHD4 recruited by TBX5 activates and represses the expression of hundreds of genes in aCMs, including genes selectively expressed in aCMs. The increase in AF vulnerability of CHD4 mice suggested that some of these genes may be associated with the maintenance of atrial rhythm. Thus, we compared genes that are upregulated and downregulated in aCMs by inactivation of both CHD4 and TBX5 with a list of genes associated with human AF by GWAS or familial studies (**Fig. 7a**). We found 42 candidate genes, including *MYL4*, which is enhanced by CHD4 and TBX5 expression, and *FBN2*, which was repressed by CHD4 and TBX5 (**Fig. 7b**).

**Fig. 7.**
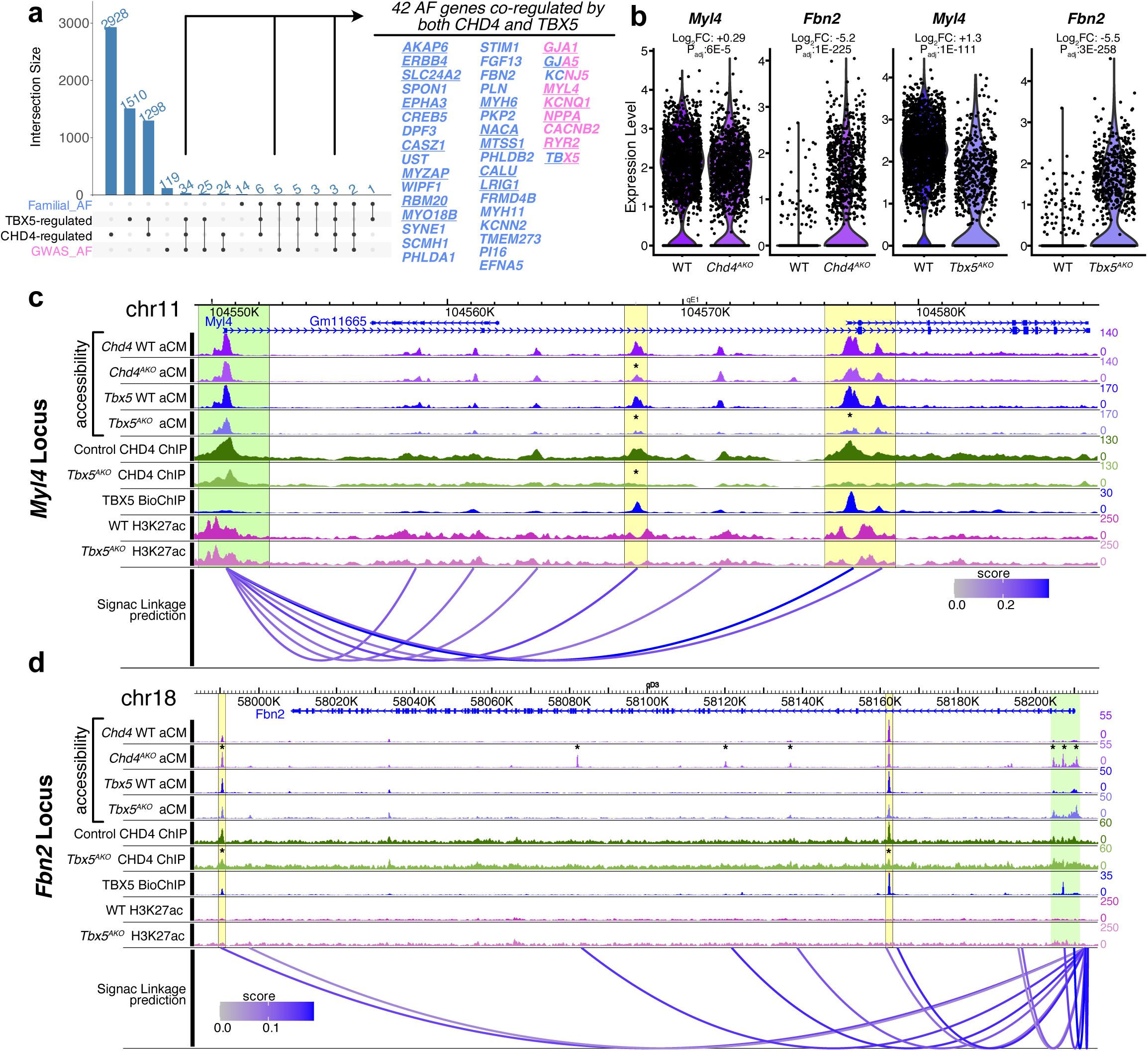
Atrial fibrillation genes are co-regulated by CHD4 and TBX5. **a**, Intersection of genes regulated by CHD4 or TBX5 (**Supp. Fig. 2a**) with genes that increase AF risk in GWAS (blue) or familial (pink) studies. CHD4 and TBX5 co-regulated 42 genes associated with AF. Underlined genes were upregulated by CHD4 and TBX5. The remainder were downregulated by CHD4 and TBX5. **b**, Expression changes for *Myl4* and *Fbn2* in *Tbx5^AKO^* and *Chd4^AKO^*snRNA-seq datasets. Log FC and Bonferroni-adjusted Wilcoxon rank-sum P values are shown. **c,d**, Visualization of chromatin features at the *Myl4* and *Fbn2* loci. Enhancer-gene linkage predictions were calculated based on the correlation between gene expression and locus accessibility in each cell from the CHD4 multiomics experiment. ATAC data from WT and *Tbx5^AKO^* aCMs were from GSE222970. TBX5 BioChIP data were from GSE215065. H3K27ac data were from GSE222970.

To explain *Myl4* is activation by CHD4 and TBX5, we annotated the *Myl4* locus with: (1) chromatin accessibility from WT and *Chd4^AKO^* aCMs from the snATACseq experiment; (2) chromatin accessibility from WT and *Tbx5^AKO^* aCMs from the snATACseq experiment^6^; (3) CHD4 ChIP-seq from WT and *Tbx5^AKO^* aCMs^6^; (4) TBX5 bioChIP-seq from WT aCMs^2^; (5) H3K27ac ChIP-seq from WT and *Tbx5^AKO^* aCMs^6^; and (6) enhancer-gene linkage predictions generated by examining the correlation between gene expression and the accessibility of in each cell from the CHD4 multiomics experiment (**Fig. 7c**). *Myl4* contains two predicted enhancers with TBX5 and CHD4 occupancy, chromatin accessibility, and flanking H3K27ac signal (yellow highlight). For the more 5’ region, inactivating TBX5 results in the loss of CHD4 occupancy, chromatin accessibility, and flanking H3K27ac signal. Therefore, TBX5 recruits CHD4 and both factors promote the accessibility of this internal *Myl4* enhancer region to promote its expression. The *Myl4* promoter (green highlight) and more 3’ enhancer region are affected similarly following TBX5 inactivation; however, the loss of accessibility in CHD4 KO aCMs was not significant, perhaps explaining the greater magnitude of *Myl4* downregulation in TBX5 KO aCMs compared with CHD4 KO aCMs.

In contrast to *Myl4, Fbn2* expression was upregulated by either CHD4 or TBX5 inactivation. Interrogating the *Fbn2* locus in the same fashion revealed that TBX5 and CHD4 each occupied an intronic and an intergenic accessible region, which was accessible but which lacked H3K27ac (yellow highlights, **Fig. 7d**). Although these regions contact the *Fbn2* promoter, the promoter had low accessibility (green highlight). TBX5 inactivation resulted in loss of CHD4 occupancy and decreased chromatin accessibility at the distal region. However, the promoter accessibility increased, which correlated with *Fbn2* upregulation. CHD4 inactivation had mixed effects on distal region accessibility, and increased promoter accessibility and gene expression similarly to TBX5 inactivation.

In sum, these data highlight both the activator and repressive functions of CHD4 recruited by TBX5, and they demonstrate that these two factors robustly regulate genes associated with human AF.

## Discussion

In this study, we established that CHD4 has dual activities in aCM gene regulation. First, CHD4 has a novel aCM transcriptional activator function. At 47.3% of regions where CHD4 supports genomic accessibility in WT aCMs, CHD4 occupancy required its recruitment by TBX5 (**Fig. 8, CHD4 activator role**). These regions promote aCM gene expression, and target a large percentage of genes expressed at greater levels in aCMs compared with vCMs. Second, we confirmed the canonical transcriptional repressor function of CHD4 (**Fig. 8, CHD4 repressor role**). In agreement with prior studies^11,12^, CHD4 was essential for suppressing sarcomeric genes of non-CM myocyte lineages. TBX5-dependent CHD4 recruitment accounted for a small fraction of genes repressed by CHD4 in aCMs; only 11.4% of regions that gained accessibility in *Chd4^AKO^* aCMs had CHD4 occupancy that depended on TBX5. Indeed, other cardiac TFs co-occupied a minority of *Chd4^AKO^*DARs, suggesting that CHD4 recruitment to these loci is primarily mediated by other TFs. Nonetheless, these data demonstrate that the previously described function of CHD4 in repressing non-cardiomyocyte sarcomere genes^10,12^ in fetal and ventricular CMs is active in postnatal aCMs. CHD4 canonically represses genes as a component of the NuRD complex, although some CHD4 functions are reported to be independent of the NuRD complex^32^. In future studies, it will be important to determine the protein complexes that mediate CHD4’s activating function, and why CHD4 activation occurs predominantly by TBX5-mediated recruitment. If CHD4 activation occurs by a variant of the NuRD complex, it will be essential to identify the factors that determine when the complex functions as an activator or repressor.

**Fig. 8.**
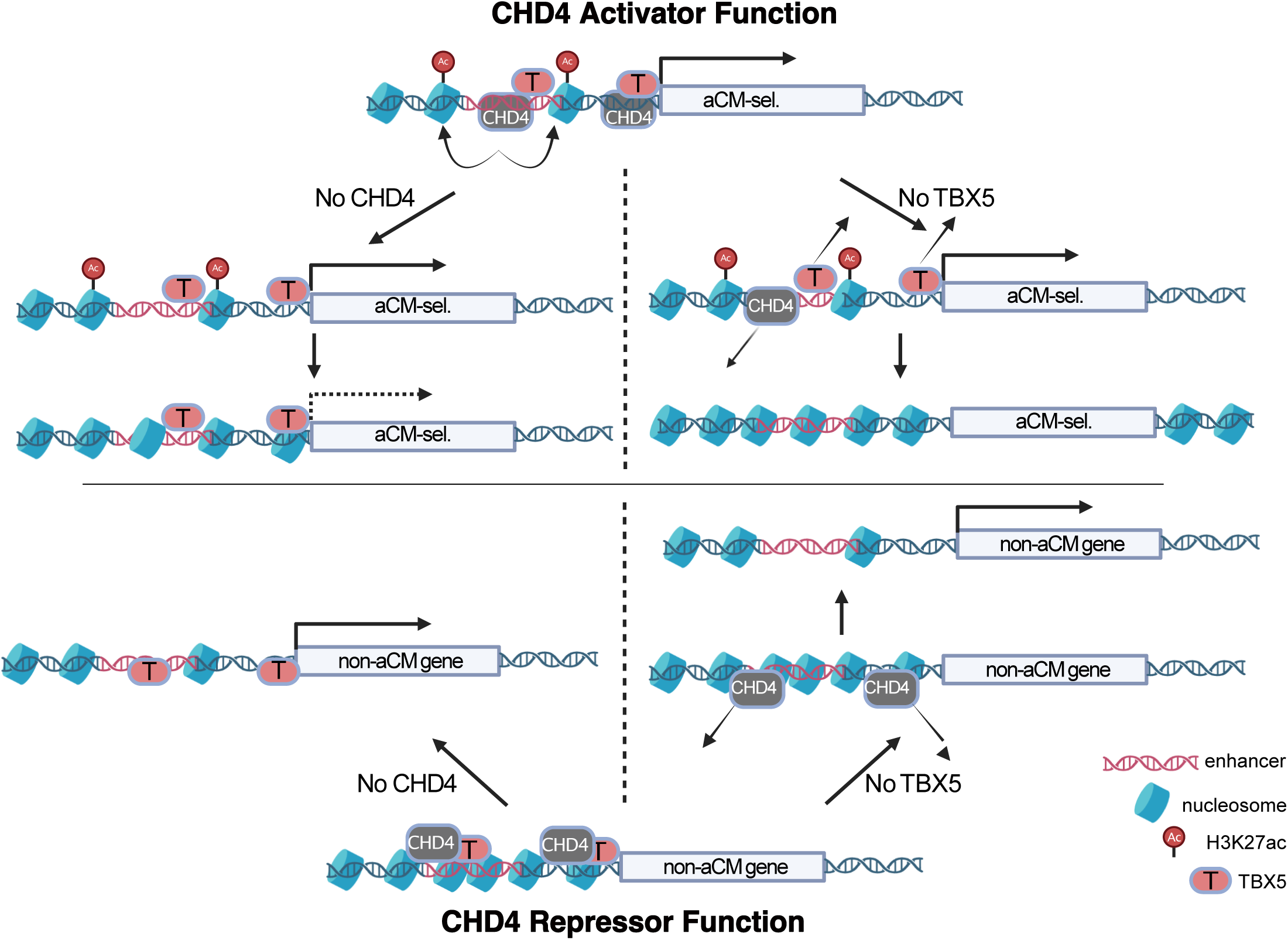
A model of TBX5-associated CHD4 repressor and activator functions. Top, CHD4 activator function. TBX5 recruits CHD4 to enhancer elements and promoters to maintain their accessibility. Inactivating CHD4 results in the loss of CHD4 from these regions, and the loss of genomic accessibility, leading to enhancer inactivation and gene downregulation. Inactivating TBX5 similarly results in the loss of CHD4, because TBX5 is necessary to recruit CHD4 to these regions. Enhancers similarly close and the gene is downregulated. Bottom, CHD4 repressor function. TBX5 recruits CHD4 to limit enhancer accessibility. The loss of CHD4 results in increased accessibility and gene activation. The loss of TBX5 reduces CHD4 recruitment, similarly causing increased accessibility and gene activation.

CHD4 inactivation in aCMs caused partially penetrant, spontaneous AF and increased vulnerability to pacing-induced AF. These results establish CHD4 as the first chromatin remodeling enzyme associated with AF. Ablation of TBX5 in postnatal aCMs causes rapid development of spontaneous AF due to loss of aCM identity^6^ and misexpression of Ca^2+^ handling genes^15,33^. Here we show that CHD4 collaborates with TBX5 to maintain atrial enhancer accessibility and activate expression of genes selectively expressed in aCMs. Human genetic studies identified over 200 genes associated with increased AF risk^34–36^, and 95 of these genes are regulated by TBX5 (26), CHD4 (27), or both (42). These data suggest that increased AF vulnerability in *Chd4^AKO^* mice is likely due to dysregulation of multiple genes. Of the AF risk genes co-regulated by CHD4 and TBX5, nine (*Tbx5, Mtss1, Myl4, Kcnq1, Nppa, Dpf3, Gja1, Myo18b,* and *Casz1*) had all of the hallmarks of being activated by the novel TBX5-dependent CHD4 gene activation mechanism: co-occupied by TBX5-enhanced CHD4 binding, loss of accessibility with loss of CHD4, and downregulation with both loss of TBX5 and loss of CHD4. TBX5-dependent CHD4 gene activation of these AF risk genes highlights the significance of the CHD4 activating mechanism to aCM rhythm homeostasis. That TBX5-CHD4 may activate *Tbx5* itself suggests an intriguing potential feed-forward loop that reinforces aCM identity. This feed-forward circuit was suggested by the downregulation of *Tbx5* in *Chd4^AKO^*.

In patients, AF usually starts as paroxysmal AF and progresses to sustained AF^37^. In experimental models brief episodes of pacing-induced AF become progressively longer until sustained AF develops^38^. The mechanisms by which episodes of AF remodel the atria’s electrical properties to facilitate and sustain AF are unclear, but the persistent effect of prior episodes of AF on atrial properties suggests an important function for epigenetic alterations in the atrial gene program. Histone deacetylases participate in AF pathogenesis^39^, and histone deacetylase inhibitors reduce AF vulnerability in experimental models^40^. As an important epigenetic regulator that interacts with histone deacetylases as well as key TFs that govern aCM identity and atrial rhythm homeostasis, CHD4 is well positioned to be a central epigenetic factor that participates in the transcriptional reprogramming that accompanies AF progression. Additional studies are needed to test this intriguing hypothesis.

## Methods

### Mice

All animal research was conducted under protocols approved by the Institutional Animal Care and Use Committees (IACUC) at Boston Children’s Hospital or the University of North Carolina, Chapel Hill. The following mouse strains were described and used in this study: *Tbx5^Flox^* ^20^, *Brg1^Flox^*^13^, and *Chd4^Flox^* ^41^. All mouse strains were maintained on a mixed genetic background, with approximately equal numbers of male and female mice used in each experiment.

### Echocardiography

Echocardiography was conducted using a Vevo 3100 imaging system (Visual Sonics) with awake animals positioned in a standard hand grip. Cardiac function was assessed from the parasternal short axis view. Image acquisition and analysis was performed blinded to group assignment.

### Surface and intracardiac electrocardiograms and atrial stimulation

Mice were anesthetized with 3% isoflurane and maintained at 37°C on a heating pad throughout the experiment. They were positioned in dorsal recumbency, and platinum electrodes were inserted subcutaneously on each of the four limbs. ECG recordings were collected for 3 minutes using the Iworks IX-ECG6 recorder and LabScribe software. For each mouse, 1500 beats were analyzed to calculate the standard deviation of the RR interval (SDRR). RR intervals were exported from LabScribe 3 software, and SDRR was calculated in Excel. All ECGs were recorded with blinding to treatment groups.

Programmed atrial stimulation protocols were performed under isoflurane anesthesia. An octapolar electrophysiology catheter (ADInstruments) was inserted through a right jugular cutdown. Programmed right atrial stimulation was performed using S1 drive trains of 80 ms intervals for 20 beats, followed by five extra stimulations at 50 ms intervals each. The interval of the five extra stimulations was progressively decreased by 5 ms per round of stimulation from 50 ms to 20 ms. Data acquisition and analysis were performed blinded to group assignment. An animal was classified as having AF if at least three seconds of AF was observed following one of the pacing rounds.

### AAV purification

AAVs were purified and quantified using a standard protocol. Briefly, AAV was produced in HEK293T cells (ATCC CRL-3216) with AAV9 Rep/Cap and purified via iodixanol density gradient ultracentrifugation as described^42^. Viral concentration was determined by qPCR. In all experiments, AAV (2 × 10^11^ viral genomes per gram body weight) was administered subcutaneously to neonatal mice.

### Histological staining and immunostaining

Mouse hearts were harvested, rinsed with ice-cold 1× PBS, fixed in 4% PFA, and then washed three times in 1× PBS for 10 minutes. Hearts were then processed through a standard ethanol dehydration gradient and embedded in paraffin. Sections of 7µm were cut, floated in a water bath (40°C), and attached to charged slides. Antigen retrieval was performed using citrate buffer (pH=6.0) and the 2100 retriever system (Aptum Biologics). Following antigen retrieval, sections were permeabilized (PBS+0.1% triton X 100) for 20 minutes, washed 2x in PBS, and blocked. Primary antibody was incubated overnight (antibody information and concentrations are available in **supplemental Table 7**) at 4°C. Slides were washed 3x with PBS, and secondary antibodies were applied (Donkey anti-Rabbit Alexa Fluor 488, Life Technologies A32790, or Donkey anti-Mouse Alexa Fluor 555, Life technologies A31570, at concentrations of 1:300) with Alexa Fluor 647 linked Wheat Germ Agglutinin (Molecular Probes, W32466, 1:500) at RT for 1 hour. Slides were washed 4x, coverslipped using VECTASHIELD mounting medium containing DAPI (VWR 101098-044), and sealed with fingernail polish. Information regarding primary antibodies and concentrations used is in **Supp. Table 7**.

### Microscopy

Confocal images were acquired on an Olympus FV3000RS confocal microscope with a 60x oil immersion lens (NA = 1.4). For fibrosis measurement, mouse hearts were excised, fixed in 4% PFA overnight, dehydrated in an ethanol gradient, embedded in paraffin, and sectioned at 7 µm. Sections were then dewaxed, rehydrated, and stained using Picrosirius red/fast green staining to visualize fibrosis. A Keyence widefield microscope was used to make a tiled image of the section using a 20x objective. Using FIJI or ImageJ, the image was split into separate channels and the green channel was used to maximize contrast. With the adjust threshold tool, the slider was adjusted to highlight the fibrotic tissue, and the number of selected pixels was recorded. The slider was then adjusted to highlight the entire tissue area without including the background, and the number of selected pixels was recorded. The percent of fibrotic tissue was calculated by dividing the fibrotic tissue pixels by the total tissue pixels and multiplying by 100.

### Combined snRNA and snATAC sequencing

Left and right atrial samples were flash-frozen in liquid nitrogen and stored at −80°C. To minimize variability, samples were collected and processed in two rounds, with Gel Bead-In Emulsions (GEMs) encapsulated in two rounds. Nuclei were isolated following a published protocol with minor modifications^43^. Briefly, frozen atria were resuspended in homogenization buffer (250 mM sucrose, 25 mM KCl, 5 mM MgCl₂, 10 mM Tris, pH 8, 1 µM DTT, with protease inhibitor, 0.4 U/µl RNaseIn, 0.2 U/µl SuperaseIn, and 0.1% Triton X-100). The samples were homogenized with a Qiagen TissueLyser II at 25 Hz for 90 seconds using a 1 mm ball bearing, then filtered through a 40 µm pluriselect mini-strainer, centrifuged at 500g for 5 minutes, and resuspended in storage buffer (4% BSA in PBS, 0.2 U/µl RNaseIn). Samples were stained with 7-aminoactinomycin D at a final concentration of 1 µg/ml and purified by FACS sorting. For each multiome sample, 300,000 nuclei were collected into storage buffer. The nuclei were then gently permeabilized with multiome lysis buffer (10 mM Tris-HCl pH 7.4, 10 mM NaCl, 3 mM MgCl₂, 0.1% Tween-20, 0.1% NP-40, 0.01% Digitonin, 1% BSA, 1 mM DTT, 1 U/µl RNaseIN), washed twice with wash buffer (10 mM Tris-HCl pH 7.4, 10 mM NaCl, 3 mM MgCl₂, 1% BSA, 0.1% Tween-20, 1 mM DTT, 1 U/µl RNaseIN), and resuspended in 1x nuclei buffer (10x Genomics). Nuclei concentration was determined using the Countess 3 system, and nuclei were then processed with the 10x Genomics Next GEM Single Cell Multiome ATAC + Gene Expression Reagents kit. The resulting snRNA and snATAC libraries were quality-checked with a TapeStation (Agilent) and sequenced on an Illumina NovaSeq instrument. Sequencing metrics are summarized in **Supp. Table 8**.

### ChIP-seq

Atria from control or *Tbx5^AKO^* mice were harvested and nuclei were extracted following the same protocol as for multiomics. Nuclei from aCMs were enriched by purification by MACS for PCM1, as described^44^. Formaldehyde was added to the nuclei for a final concentration of 2%, and the nuclei were fixed with gentle rocking at room temperature for 15 minutes. The reaction was quenched with glycine, and the nuclei were pelleted and counted. In total, the number of input nuclei for each ChIP reaction was approximately 6.5 million. Nuclei pellets were snap frozen in liquid nitrogen for storage. Briefly, nuclei pellets were resuspended and washed once with the Hypotonic buffer (10 mM HEPES-NaOH pH7.9, 10mM KCl, 1.5mM MgCl_2_, 340mM sucrose, 10% glycerol, 0.1% Triton-X 100, protease inhibitors) followed by spinning down at 2000 g at 4°C for 5 min, and then the pellets were resuspended and dounced 20 times with a tissue grinder on ice in shearing buffer (10 mM Tris-HCl pH8.0, 1 mM EDTA, 0.5 mM EGTA, 0.5 mM PMSF, 5 mM NaBut, 0.1% SDS, protease inhibitors). Homogenates were transferred and sonicated in a Covaris E220 Focused-ultrasonicator with settings of power peak 175, duty factor 10, 200 counts for 1200 seconds at 4°C. Following sonication, samples were centrifuged at 20000 g at 4°C for 10 min. 50 µg of chromatin was used as an input for each sample and was diluted 1:1 with 2×IP buffer (20 mM Tris-HCl pH8.0, 300 mM NaCl, 2 mM EDTA, 20% glycerol, 1% Triton-X 100, 5 mM NaBut, 0.5 mM PMSF, protease inhibitors). Ten percent of chromatin from each sample was used as an input control and stored at −20°C until the reverse-crosslink step. CHD4 was immunoprecipitated using a specific antibody (Abcam ab70469) at a concentration of 5 µg per ChIP reaction, rotating overnight at 4°C.. 20 μl of each Protein A (Thermo, 10001D) and Protein G Dynabeads (Thermo, 10003D) were blocked overnight at 4°C with 75 mg/ml bovine serum albumin (final concentration: 5 mg/ml, Sigma, A7906). The next day, blocked beads were mixed with the chromatin and rotated for 3 hrs at 4°C, then washed sequentially with low salt buffer (20 mM Tris-HCl pH8.0, 150 mM NaCl, 2 mM EDTA, 1% Triton-X 100, 0.1% SDS), high salt buffer (20 mM Tris-HCl pH8.0, 300 mM NaCl, 2 mM EDTA, 1% Triton-X 100, 0.1% SDS), LiCl buffer (20 mM Tris-HCl pH8.0, 250 mM LiCl, 2 mM EDTA, 1% NP-40, 1%/vol NaDeoxycholate), and TE10/1 buffer (10 mM Tris-HCl pH8.0, 1 mM EDTA). Chromatin was eluted with freshly prepared elution buffer (1% SDS, 0.1 M NaHCO_3_, 5 mM DTT) for 1 hr at 65°C. Eluted chromatin and the input control samples were incubated with 6 µl of 5 M NaCl and 0.5 µl of RNase cocktail for 4 hrs at 65°C to reverse crosslink, then 2 µl of 1 M Tris-HCl pH 6.8 and 2 µl of 20 mg/ml Proteinase K were added and incubated for 1 hr at 65°C. Enriched and eluted DNA was then purified using Ampure XP beads (Beckman Coulter, A63880). Libraries were prepared using the ThruPLEX DNA-seq kit (TaKaRa), and sequenced on a NovaSeq-S1. Paired-end sequencing was performed (2x 50nt). Following sequencing, genomic alignment of FASTQ files was performed with bowtie2 to the mm10 genome, and the aligned .bam files were filtered to remove reads with affinity scores less than 20[1] _[WS2]_ and to remove mitochondrial reads. Peak calling was performed using MACS2, by comparing each sample to their input sample, using a q-value<0.05. Enriched peaks from biological replicates were combined, and differential peaks were determined by comparing the combined ChIP control data with the combined ChIP TBX5 KO data using R package DiffBind^16^. Peak annotation was performed by using R package ChIPseeker^45^.

### Bulk RNA-seq

Both left and right atria were collected and combined as one biological replicate for each heart. The tissues were homogenized in ice-cold TRIzol reagent with a POLYTRON homogenizer, followed by adding 0.2 mL chloroform. The total RNA was extracted using the PureLink RNA Mini Kit (ThermoFisher, 12183018A). DNA was removed by incubating the eluate with PureLink DNase (ThermoFisher, 12185010) at room temperature for 15 min. The concentration and quality of purified RNA were assessed by Tape Station (Agilent). RNA integrity numbers (RIN) ranged from 8.0 to 10.0. Purified poly-A RNA that had undergone two rounds of oligo-dT selection was converted into cDNA and used to generate RNA-seq libraries. Libraries were sequenced (75-nt paired-end reads; Illumina HiSeq 2500) to a target depth of >30 million reads. Reads were aligned to the mm10 reference genome using STAR via the bcbio-nextgen RNA sequencing pipeline. RNAseq analysis was performed using R package DESeq2.

### Single cell data analysis and object creation

Data outputs from the 10x Cellranger-ARC (version 2.0.1) software package were processed using the Seurat (version 5.1.0) and Signac (version 1.13.0) packages in R^46,47^ (version 4.4.1). In brief, Cellranger-ARC was run on each sample to generate output files, which were used to construct Seurat objects. For snATAC-seq data analysis, a shared peak set was created by integrating peaks identified across samples, following the Signac peak merging vignette. Quality control (QC) was applied to each Seurat object using consistent metrics: 250 < nCount_ATAC < 100,000; 250 < nCount_RNA < 35,000; nucleosome_signal < 2; TSS.enrichment > 1; and percent mitochondrial reads < 25. Doublets were detected and removed using DoubletFinder^48^ on the RNA portion of each sample. After QC, individual objects were merged. Final QC metrics for each individual sample after filtering are available in **Supp. Fig. 4**.

### Object creation and cluster identification

Data normalization and scaling were performed for the RNA dataset using Seurat SCTransform() to account for sequencing depth and other sources of variation. Then principle component analysis (dims 1:20) was performed. An RNA UMAP was created using the 30 nearest neighbors and the first 20 PCs. For the ATAC dataset, Term frequency-inverse document frequency (TF-IDF) normalization was applied. All peaks were retained for top feature selection, and Singular Value Decomposition (SVD) was performed on the top ATAC features to create a low dimensional representation based on latent semantic indexing (LSI). A UMAP embedding was created from the LSI-reduced data, using dimensions 2-15 for downstream analysis (dimension 1 was excluded because it usually corresponds to changes in sequencing depth). To integrate the RNA and ATAC data and produce a WNN UMAP embedding, the Seurat Findmultimodalneighbors() function was performed using RNA PCA 1:15 and ATAC LSI 2:15. Finally, cell clustering was performed by applying the Louvain algorithm to the WNN embedding at a resolution of 0.8. Broad cluster identification was performed using the SCtype package^26^ with the ‘Heart’ encyclopedia. SCtype annotations were then refined using three previously annotated top marker genes for different cell types found in the atria^6^. Differential gene expression and differential genomic accessibility analyses were performed comparing clusters using the FindMarkers function on either the RNA or the ATAC assays. Differential accessibility was performed by logistic regression using Signac, with thresholds of |log_2_FC| > 0.25 and P_Adj_ <0.05. The same thresholds were used to identify differentially expressed genes. Differential gene expression analysis was performed using the Wilcoxon test. Genes neighboring enhancer regions were identified using the ClosestFeature command in Signac. To generate bigwig files from the ATAC data for individual clusters, fragment files for each of the samples were split using the list of cell barcodes composing the cluster of interest using the sinto package. The bamcoverage tool was used to create RPGC normalized .bigwig files. Region-gene linkage scores for enhancer prediction were determined by identifying correlations between the accessibility of different genomic regions with the expression of the gene. This analysis was constrained to genomic regions +/−500 kb from the TSS of the examined gene^49^.

### Statistics and Reproducibility

Results were expressed as mean ± SEM. Statistical tests, indicated in figure legends, were performed using Graphpad Prism 9 or R. P<0.05 was used as the statistical threshold for significance. Micrographs are representative of three independent experiments (unless indicated otherwise). Datapoints in graphs represent unique biological replicates.

## Data Availability

New data generated by the study will be deposited and publicly accessible at the time of publication. Other datasets were accessed from GSE215065^2^ and GSE222970^6^. All other data supporting the findings in this study are included in the main article and associated files. Source data are provided with this manuscript.

## Acknowledgements

MES was supported by K99HL173573. WS was supported by the Center for Heart and Vascular Research COBRE (P20GM152326). WTP was supported by R01HL156503. The funders had no role in the study design, data collection and analysis, decision to publish or preparation of the manuscript.

## References

1. Zhou, P., VanDusen, N. J., Zhang, Y., Cao, Y., Sethi, I., Hu, R., Zhang, S., Wang, G., Ye, L., Mazumdar, N., Chen, J., Zhang, X., Guo, Y., Li, B., Ma, Q., Lee, J. Y., Gu, W., Yuan, G.-C., Ren, B., Chen, K. & Pu, W. T. Dynamic changes in P300 enhancers and enhancer-promoter contacts control mouse cardiomyocyte maturation. Dev. Cell 58, 898–914.e7 (2023).

2. Cao, Y., Zhang, X., Akerberg, B. N., Yuan, H., Sakamoto, T., Xiao, F., VanDusen, N. J., Zhou, P., Sweat, M. E., Wang, Y., Prondzynski, M., Chen, J., Zhang, Y., Wang, P., Kelly, D. P. & Pu, W. T. In Vivo Dissection of Chamber-Selective Enhancers Reveals Estrogen-Related Receptor as a Regulator of Ventricular Cardiomyocyte Identity. Circulation 147, 881–896 (2023).

3. Kathiriya, I. S., Dominguez, M. H., Rao, K. S., Muncie-Vasic, J. M., Devine, W. P., Hu, K. M., Hota, S. K., Garay, B. I., Quintero, D., Goyal, P., Matthews, M. N., Thomas, R., Sukonnik, T., Miguel-Perez, D., Winchester, S., Brower, E. F., Forjaz, A., Wu, P.-H., Wirtz, D., Kiemen, A. L. & Bruneau, B. G. A disrupted compartment boundary underlies abnormal cardiac patterning and congenital heart defects. bioRxivorg 2024.02.05.578995 (2024).

4. Bruneau, B. G., Logan, M., Davis, N., Levi, T., Tabin, C. J., Seidman, J. G. & Seidman, C. E. Chamber-specific cardiac expression of Tbx5 and heart defects in Holt-Oram syndrome. Dev. Biol. 211, 100–108 (1999).

5. Moskowitz, I. P. G., Pizard, A., Patel, V. V., Bruneau, B. G., Kim, J. B., Kupershmidt, S., Roden, D., Berul, C. I., Seidman, C. E. & Seidman, J. G. The T-Box transcription factor Tbx5 is required for the patterning and maturation of the murine cardiac conduction system. Development 131, 4107–4116 (2004).

6. Sweat, M. E., Cao, Y., Zhang, X., Burnicka-Turek, O., Perez-Cervantes, C., Arulsamy, K., Lu, F., Keating, E. M., Akerberg, B. N., Ma, Q., Wakimoto, H., Gorham, J. M., Hill, L. D., Kyoung Song, M., Trembley, M. A., Wang, P., Gianeselli, M., Prondzynski, M., Bortolin, R. H., Bezzerides, V. J., Chen, K., Seidman, J. G., Seidman, C. E., Moskowitz, I. P. & Pu, W. T. Tbx5 maintains atrial identity in post-natal cardiomyocytes by regulating an atrial-specific enhancer network. Nat Cardiovasc Res 2, 881–898 (2023).

7. Cirillo, L. A., Lin, F. R., Cuesta, I., Friedman, D., Jarnik, M. & Zaret, K. S. Opening of compacted chromatin by early developmental transcription factors HNF3 (FoxA) and GATA-4. Mol. Cell 9, 279–289 (2002).

8. Takeuchi, J. K., Lou, X., Alexander, J. M., Sugizaki, H., Delgado-Olguín, P., Holloway, A. K., Mori, A. D., Wylie, J. N., Munson, C., Zhu, Y., Zhou, Y.-Q., Yeh, R.-F., Henkelman, R. M., Harvey, R. P., Metzger, D., Chambon, P., Stainier, D. Y. R., Pollard, K. S., Scott, I. C. & Bruneau, B. G. Chromatin remodelling complex dosage modulates transcription factor function in heart development. Nat. Commun. 2, 187 (2011).

9. Waldron, L., Steimle, J. D., Greco, T. M., Gomez, N. C., Dorr, K. M., Kweon, J., Temple, B., Yang, X. H., Wilczewski, C. M., Davis, I. J., Cristea, I. M., Moskowitz, I. P. & Conlon, F. L. The Cardiac TBX5 Interactome Reveals a Chromatin Remodeling Network Essential for Cardiac Septation. Dev. Cell 36, 262–275 (2016).

10. Wilczewski, C. M., Hepperla, A. J., Shimbo, T., Wasson, L., Robbe, Z. L., Davis, I. J., Wade, P. A. & Conlon, F. L. CHD4 and the NuRD complex directly control cardiac sarcomere formation. Proc. Natl. Acad. Sci. U. S. A. 115, 6727–6732 (2018).

11. Robbe, Z. L., Shi, W., Wasson, L. K., Scialdone, A. P., Wilczewski, C. M., Sheng, X., Hepperla, A. J., Akerberg, B. N., Pu, W. T., Cristea, I. M., Davis, I. J. & Conlon, F. L. CHD4 is recruited by GATA4 and NKX2-5 to repress noncardiac gene programs in the developing heart. Genes Dev. 36, 468–482 (2022).

12. Gómez-Del Arco, P., Perdiguero, E., Yunes-Leites, P. S., Acín-Pérez, R., Zeini, M., Garcia-Gomez, A., Sreenivasan, K., Jiménez-Alcázar, M., Segalés, J., López-Maderuelo, D., Ornés, B., Jiménez-Borreguero, L. J., D’Amato, G., Enshell-Seijffers, D., Morgan, B., Georgopoulos, K., Islam, A. B. M. M. K., Braun, T., de la Pompa, J. L., Kim, J., Enriquez, J. A., Ballestar, E., Muñoz-Cánoves, P. & Redondo, J. M. The Chromatin Remodeling Complex Chd4/NuRD Controls Striated Muscle Identity and Metabolic Homeostasis. Cell Metab. 23, 881–892 (2016).

13. Sumi-Ichinose, C., Ichinose, H., Metzger, D. & Chambon, P. SNF2beta-BRG1 is essential for the viability of F9 murine embryonal carcinoma cells. Mol. Cell. Biol. 17, 5976–5986 (1997).

14. Hulsurkar, M. M., Lahiri, S. K., Moore, O., Moreira, L. M., Abu-Taha, I., Kamler, M., Dobrev, D., Nattel, S., Reilly, S. & Wehrens, X. H. T. Atrial-Specific LKB1 Knockdown Represents a Novel Mouse Model of Atrial Cardiomyopathy With Spontaneous Atrial Fibrillation. Circulation 144, 909–912 (2021).

15. Nadadur, R. D., Broman, M. T., Boukens, B., Mazurek, S. R., Yang, X., van den Boogaard, M., Bekeny, J., Gadek, M., Ward, T., Zhang, M., Qiao, Y., Martin, J. F., Seidman, C. E., Seidman, J., Christoffels, V., Efimov, I. R., McNally, E. M., Weber, C. R. & Moskowitz, I. P. Pitx2 modulates a Tbx5-dependent gene regulatory network to maintain atrial rhythm. Sci. Transl. Med. 8, 354ra115 (2016).

16. Stark, R., Brown, G. & Others. DiffBind: differential binding analysis of ChIP-Seq peak data. R package version 100, (2011).

17. Yangpo Cao, Xiaoran Zhang, Brynn Akerberg, Haiyun Yuan, Tomoya Sakamoto, Feng Xiao, Nathan VanDusen, Pingzhu Zhou, Mason Sweat, Yi Wang, Maksymilian Prondzynski, Jian Chen, Yan Zhang, Peizhe Wang, Daniel Kelly, William Pu. In vivo dissection of chamber selective enhancers reveals estrogen-related receptor as a regulator of ventricular cardiomyocyte identity. Circulation

18. Akerberg, B. N., Gu, F., VanDusen, N. J., Zhang, X., Dong, R., Li, K., Zhang, B., Zhou, B., Sethi, I., Ma, Q., Wasson, L., Wen, T., Liu, J., Dong, K., Conlon, F. L., Zhou, J., Yuan, G.-C., Zhou, P. & Pu, W. T. A reference map of murine cardiac transcription factor chromatin occupancy identifies dynamic and conserved enhancers. Nat. Commun. 10, 4907 (2019).

19. Shi, W., Wasson, L. K., Dorr, K. M., Robbe, Z. L., Wilczewski, C. M., Hepperla, A. J., Davis, I. J., Seidman, C. E., Seidman, J. G. & Conlon, F. L. CHD4 and SMYD1 repress common transcriptional programs in the developing heart. Development 151, dev202505 (2024).

20. Bruneau, B. G., Nemer, G., Schmitt, J. P., Charron, F., Robitaille, L., Caron, S., Conner, D. A., Gessler, M., Nemer, M., Seidman, C. E. & Seidman, J. G. A murine model of Holt-Oram syndrome defines roles of the T-box transcription factor Tbx5 in cardiogenesis and disease. Cell 106, 709–721 (2001).

21. Misra, C., Chang, S.-W., Basu, M., Huang, N. & Garg, V. Disruption of myocardial Gata4 and Tbx5 results in defects in cardiomyocyte proliferation and atrioventricular septation. Hum. Mol. Genet. 23, 5025–5035 (2014).

22. Ghosh, T. K., Song, F. F., Packham, E. A., Buxton, S., Robinson, T. E., Ronksley, J., Self, T., Bonser, A. J. & Brook, J. D. Physical interaction between TBX5 and MEF2C is required for early heart development. Mol. Cell. Biol. 29, 2205–2218 (2009).

23. Lu, F., Ma, Q., Xie, W., Liou, C. L., Zhang, D., Sweat, M. E., Jardin, B. D., Naya, F. J., Guo, Y., Cheng, H. & Pu, W. T. CMYA5 establishes cardiac dyad architecture and positioning. Nat. Commun. 13, 2185 (2022).

24. Hulsmans, M., Schloss, M. J., Lee, I.-H., Bapat, A., Iwamoto, Y., Vinegoni, C., Paccalet, A., Yamazoe, M., Grune, J., Pabel, S., Momin, N., Seung, H., Kumowski, N., Pulous, F. E., Keller, D., Bening, C., Green, U., Lennerz, J. K., Mitchell, R. N., Lewis, A., Casadei, B., Iborra-Egea, O., Bayes-Genis, A., Sossalla, S., Ong, C. S., Pierson, R. N., Aster, J. C., Rohde, D., Wojtkiewicz, G. R., Weissleder, R., Swirski, F. K., Tellides, G., Tolis, G., Jr, Melnitchouk, S., Milan, D. J., Ellinor, P. T., Naxerova, K. & Nahrendorf, M. Recruited macrophages elicit atrial fibrillation. Science 381, 231–239 (2023).

25. Laforest, B., Dai, W., Tyan, L., Lazarevic, S., Shen, K. M., Gadek, M., Broman, M. T., Weber, C. R. & Moskowitz, I. P. Atrial fibrillation risk loci interact to modulate Ca2+-dependent atrial rhythm homeostasis. J. Clin. Invest. 129, 4937–4950 (2019).

26. Ianevski, A., Giri, A. K. & Aittokallio, T. Fully-automated and ultra-fast cell-type identification using specific marker combinations from single-cell transcriptomic data. Nat. Commun. 13, 1246 (2022).

27. GTEx Consortium. The Genotype-Tissue Expression (GTEx) project. Nat. Genet. 45, 580–585 (2013).

28. Lai, A. Y. & Wade, P. A. Cancer biology and NuRD: a multifaceted chromatin remodelling complex. Nat. Rev. Cancer 11, 588–596 (2011).

29. Yi, Y., Du, L., Qin, M., Chen, X.-Q., Sun, X.-N., Li, C., Du, L.-J., Liu, Y., Liu, Y., Sun, J.-Y., Tang, Z., Xu, M., Fang, B., Liu, X. & Duan, S.-Z. Regulation of atrial fibrosis by the bone. Hypertension 73, 379–389 (2019).

30. Milde-Langosch, K. The Fos family of transcription factors and their role in tumourigenesis. Eur. J. Cancer 41, 2449–2461 (2005).

31. Morita, R., Suzuki, M., Kasahara, H., Shimizu, N., Shichita, T., Sekiya, T., Kimura, A., Sasaki, K.-I., Yasukawa, H. & Yoshimura, A. ETS transcription factor ETV2 directly converts human fibroblasts into functional endothelial cells. Proc. Natl. Acad. Sci. U. S. A. 112, 160–165 (2015).

32. O’Shaughnessy, A. & Hendrich, B. CHD4 in the DNA-damage response and cell cycle progression: not so NuRDy now. Biochem. Soc. Trans. 41, 777–782 (2013).

33. Dai, W., Laforest, B., Tyan, L., Shen, K. M., Nadadur, R. D., Alvarado, F. J., Mazurek, S. R., Lazarevic, S., Gadek, M., Wang, Y., Li, Y., Valdivia, H. H., Shen, L., Broman, M. T., Moskowitz, I. P. & Weber, C. R. A calcium transport mechanism for atrial fibrillation in Tbx5-mutant mice. Elife 8, (2019).

34. Roselli, C., Chaffin, M. D., Weng, L.-C., Aeschbacher, S., Ahlberg, G., Albert, C. M., Almgren, P., Alonso, A., Anderson, C. D., Aragam, K. G., Arking, D. E., Barnard, J., Bartz, T. M., Benjamin, E. J., Bihlmeyer, N. A., Bis, J. C., Bloom, H. L., Boerwinkle, E., Bottinger, E. B., Brody, J. A., Calkins, H., Campbell, A., Cappola, T. P., Carlquist, J., Chasman, D. I., Chen, L. Y., Chen, Y.-D. I., Choi, E.-K., Choi, S. H., Christophersen, I. E., Chung, M. K., Cole, J. W., Conen, D., Cook, J., Crijns, H. J., Cutler, M. J., Damrauer, S. M., Daniels, B. R., Darbar, D., Delgado, G., Denny, J. C., Dichgans, M., Dörr, M., Dudink, E. A., Dudley, S. C., Esa, N., Esko, T., Eskola, M., Fatkin, D., Felix, S. B., Ford, I., Franco, O. H., Geelhoed, B., Grewal, R. P., Gudnason, V., Guo, X., Gupta, N., Gustafsson, S., Gutmann, R., Hamsten, A., Harris, T. B., Hayward, C., Heckbert, S. R., Hernesniemi, J., Hocking, L. J., Hofman, A., Horimoto, A. R. V. R., Huang, J., Huang, P. L., Huffman, J., Ingelsson, E., Ipek, E. G., Ito, K., Jimenez-Conde, J., Johnson, R., Jukema, J. W., Kääb, S., Kähönen, M., Kamatani, Y., Kane, J. P., Kastrati, A., Kathiresan, S., Katschnig-Winter, P., Kavousi, M., Kessler, T., Kietselaer, B. L., Kirchhof, P., Kleber, M. E., Knight, S., Krieger, J. E., Kubo, M., Launer, L. J., Laurikka, J., Lehtimäki, T., Leineweber, K., Lemaitre, R. N., Li, M., Lim, H. E., Lin, H. J., Lin, H., Lind, L., Lindgren, C. M., Lokki, M.-L., London, B., Loos, R. J. F., Low, S.-K., Lu, Y., Lyytikäinen, L.-P., Macfarlane, P. W., Magnusson, P. K., Mahajan, A., Malik, R., Mansur, A. J., Marcus, G. M., Margolin, L., Margulies, K. B., März, W., McManus, D. D., Melander, O., Mohanty, S., Montgomery, J. A., Morley, M. P., Morris, A. P., Müller-Nurasyid, M., Natale, A., Nazarian, S., Neumann, B., Newton-Cheh, C., Niemeijer, M. N., Nikus, K., Nilsson, P., Noordam, R., Oellers, H., Olesen, M. S., Orho-Melander, M., Padmanabhan, S., Pak, H.-N., Paré, G., Pedersen, N. L., Pera, J., Pereira, A., Porteous, D., Psaty, B. M., Pulit, S. L., Pullinger, C. R., Rader, D. J., Refsgaard, L., Ribasés, M., Ridker, P. M., Rienstra, M., Risch, L., Roden, D. M., Rosand, J., Rosenberg, M. A., Rost, N., Rotter, J. I., Saba, S., Sandhu, R. K., Schnabel, R. B., Schramm, K., Schunkert, H., Schurman, C., Scott, S. A., Seppälä, I., Shaffer, C., Shah, S., Shalaby, A. A., Shim, J., Shoemaker, M. B., Siland, J. E., Sinisalo, J., Sinner, M. F., Slowik, A., Smith, A. V., Smith, B. H., Smith, J. G., Smith, J. D., Smith, N. L., Soliman, E. Z., Sotoodehnia, N., Stricker, B. H., Sun, A., Sun, H., Svendsen, J. H., Tanaka, T., Tanriverdi, K., Taylor, K. D., Teder-Laving, M., Teumer, A., Thériault, S., Trompet, S., Tucker, N. R., Tveit, A., Uitterlinden, A. G., Van Der Harst, P., Van Gelder, I. C., Van Wagoner, D. R., Verweij, N., Vlachopoulou, E., Völker, U., Wang, B., Weeke, P. E., Weijs, B., Weiss, R., Weiss, S., Wells, Q. S., Wiggins, K. L., Wong, J. A., Woo, D., Worrall, B. B., Yang, P.-S., Yao, J., Yoneda, Z. T., Zeller, T., Zeng, L., Lubitz, S. A., Lunetta, K. L. & Ellinor, P. T. Multi-ethnic genome-wide association study for atrial fibrillation. Nat. Genet. 50, 1225–1233 (2018).

35. Nielsen, J. B., Thorolfsdottir, R. B., Fritsche, L. G., Zhou, W., Skov, M. W., Graham, S. E., Herron, T. J., McCarthy, S., Schmidt, E. M., Sveinbjornsson, G., Surakka, I., Mathis, M. R., Yamazaki, M., Crawford, R. D., Gabrielsen, M. E., Skogholt, A. H., Holmen, O. L., Lin, M., Wolford, B. N., Dey, R., Dalen, H., Sulem, P., Chung, J. H., Backman, J. D., Arnar, D. O., Thorsteinsdottir, U., Baras, A., O’Dushlaine, C., Holst, A. G., Wen, X., Hornsby, W., Dewey, F. E., Boehnke, M., Kheterpal, S., Mukherjee, B., Lee, S., Kang, H. M., Holm, H., Kitzman, J., Shavit, J. A., Jalife, J., Brummett, C. M., Teslovich, T. M., Carey, D. J., Gudbjartsson, D. F., Stefansson, K., Abecasis, G. R., Hveem, K. & Willer, C. J. Biobank-driven genomic discovery yields new insight into atrial fibrillation biology. Nat. Genet. 50, 1234–1239 (2018).

36. Miyazawa, K., Ito, K., Ito, M., Zou, Z., Kubota, M., Nomura, S., Matsunaga, H., Koyama, S., Ieki, H., Akiyama, M., Koike, Y., Kurosawa, R., Yoshida, H., Ozaki, K., Onouchi, Y., BioBank Japan Project, Takahashi, A., Matsuda, K., Murakami, Y., Aburatani, H., Kubo, M., Momozawa, Y., Terao, C., Oki, S., Akazawa, H., Kamatani, Y. & Komuro, I. Cross-ancestry genome-wide analysis of atrial fibrillation unveils disease biology and enables cardioembolic risk prediction. Nat. Genet. (2023). doi:10.1038/s41588-022-01284-9

37. Blum, S., Aeschbacher, S., Meyre, P., Zwimpfer, L., Reichlin, T., Beer, J. H., Ammann, P., Auricchio, A., Kobza, R., Erne, P., Moschovitis, G., Di Valentino, M., Shah, D., Schläpfer, J., Henz, S., Meyer-Zürn, C., Roten, L., Schwenkglenks, M., Sticherling, C., Kühne, M., Osswald, S., Conen, D. & Swiss-AF Investigators. Incidence and predictors of atrial fibrillation progression. J. Am. Heart Assoc. 8, e012554 (2019).

38. Wijffels, M. C., Kirchhof, C. J., Dorland, R. & Allessie, M. A. Atrial fibrillation begets atrial fibrillation. A study in awake chronically instrumented goats. Circulation 92, 1954–1968 (1995).

39. Brundel, B. J. J. M., Li, J. & Zhang, D. Role of HDACs in cardiac electropathology: Therapeutic implications for atrial fibrillation. Biochim. Biophys. Acta Mol. Cell Res. 1867, 118459 (2020).

40. Lkhagva, B., Kao, Y.-H., Chen, Y.-C., Chao, T.-F., Chen, S.-A. & Chen, Y.-J. Targeting histone deacetylases: A novel therapeutic strategy for atrial fibrillation. Eur. J. Pharmacol. 781, 250–257 (2016).

41. Williams, C. J., Naito, T., Arco, P. G.-D., Seavitt, J. R., Cashman, S. M., De Souza, B., Qi, X., Keables, P., Von Andrian, U. H. & Georgopoulos, K. The chromatin remodeler Mi-2beta is required for CD4 expression and T cell development. Immunity 20, 719–733 (2004).

42. Wang, S., Guo, Y. & Pu, W. T. AAV Gene Transfer to the Heart. Methods Mol. Biol. 2158, 269–280 (2021).

43. Nadelmann, E. R., Gorham, J. M., Reichart, D., Delaughter, D. M., Wakimoto, H., Lindberg, E. L., Litviňukova, M., Maatz, H., Curran, J. J., Ischiu Gutierrez, D., Hübner, N., Seidman, C. E. & Seidman, J. G. Isolation of Nuclei from Mammalian Cells and Tissues for Single-Nucleus Molecular Profiling. Curr Protoc 1, e132 (2021).

44. Cui, M. & Olson, E. N. Protocol for Single-Nucleus Transcriptomics of Diploid and Tetraploid Cardiomyocytes in Murine Hearts. STAR Protoc 1, 100049 (2020).

45. Yu, G., Wang, L.-G. & He, Q.-Y. ChIPseeker: an R/Bioconductor package for ChIP peak annotation, comparison and visualization. Bioinformatics 31, 2382–2383 (2015).

46. Stuart, T., Srivastava, A., Madad, S., Lareau, C. A. & Satija, R. Single-cell chromatin state analysis with Signac. Nat. Methods 18, 1333–1341 (2021).

47. Butler, A., Hoffman, P., Smibert, P., Papalexi, E. & Satija, R. Integrating single-cell transcriptomic data across different conditions, technologies, and species. Nat. Biotechnol. 36, 411–420 (2018).

48. McGinnis, C. S., Murrow, L. M. & Gartner, Z. J. DoubletFinder: Doublet Detection in Single-Cell RNA Sequencing Data Using Artificial Nearest Neighbors. Cell Syst 8, 329–337.e4 (2019).

49. Ma, S., Zhang, B., LaFave, L. M., Earl, A. S., Chiang, Z., Hu, Y., Ding, J., Brack, A., Kartha, V. K., Tay, T., Law, T., Lareau, C., Hsu, Y.-C., Regev, A. & Buenrostro, J. D. Chromatin Potential Identified by Shared Single-Cell Profiling of RNA and Chromatin. Cell 183, 1103–1116.e20 (2020).

