## Supplemental Figure 1 for "CHD4 Interacts With TBX5 to Maintain the Gene Regulatory Network of Postnatal Atrial Cardiomyocytes"

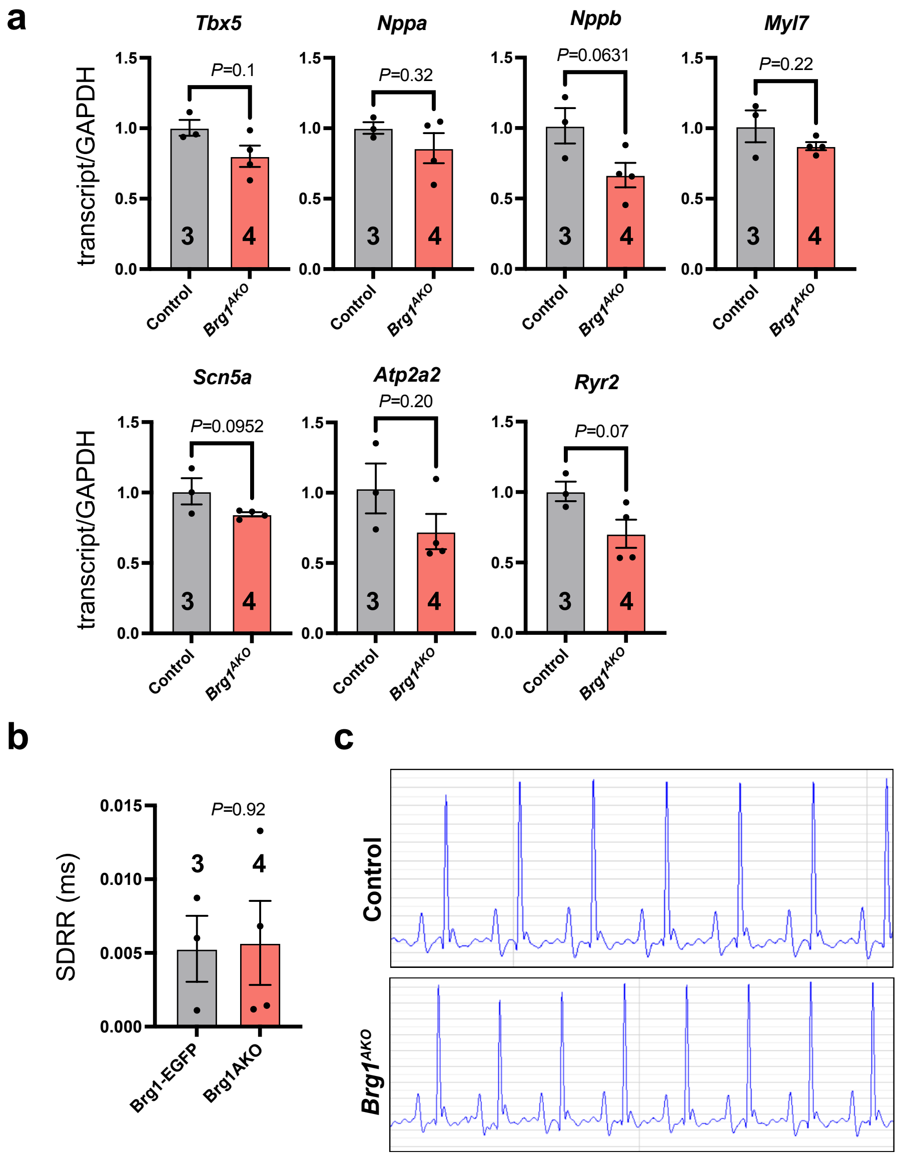


**Supp. Fig. 1. qPCR and ECG analysis of P28 *Brg1*$^{AKO}$ mice.**

**a**, RT-qPCR comparing gene expression in control and *Brg1*$^{AKO}$ atria. Two tailed *t*-test. **b**, Standard Deviation of the RR interval (SDRR) for control and *Brg1*$^{AKO}$ mice. n=3 control and 4 *Brg1*$^{AKO}$ mice, Two tailed *t*-test. **c**, Representative ECGs from control and *Brg1*$^{AKO}$ mice.
