## Supplemental Figure 2 for "CHD4 Interacts With TBX5 to Maintain the Gene Regulatory Network of Postnatal Atrial Cardiomyocytes"

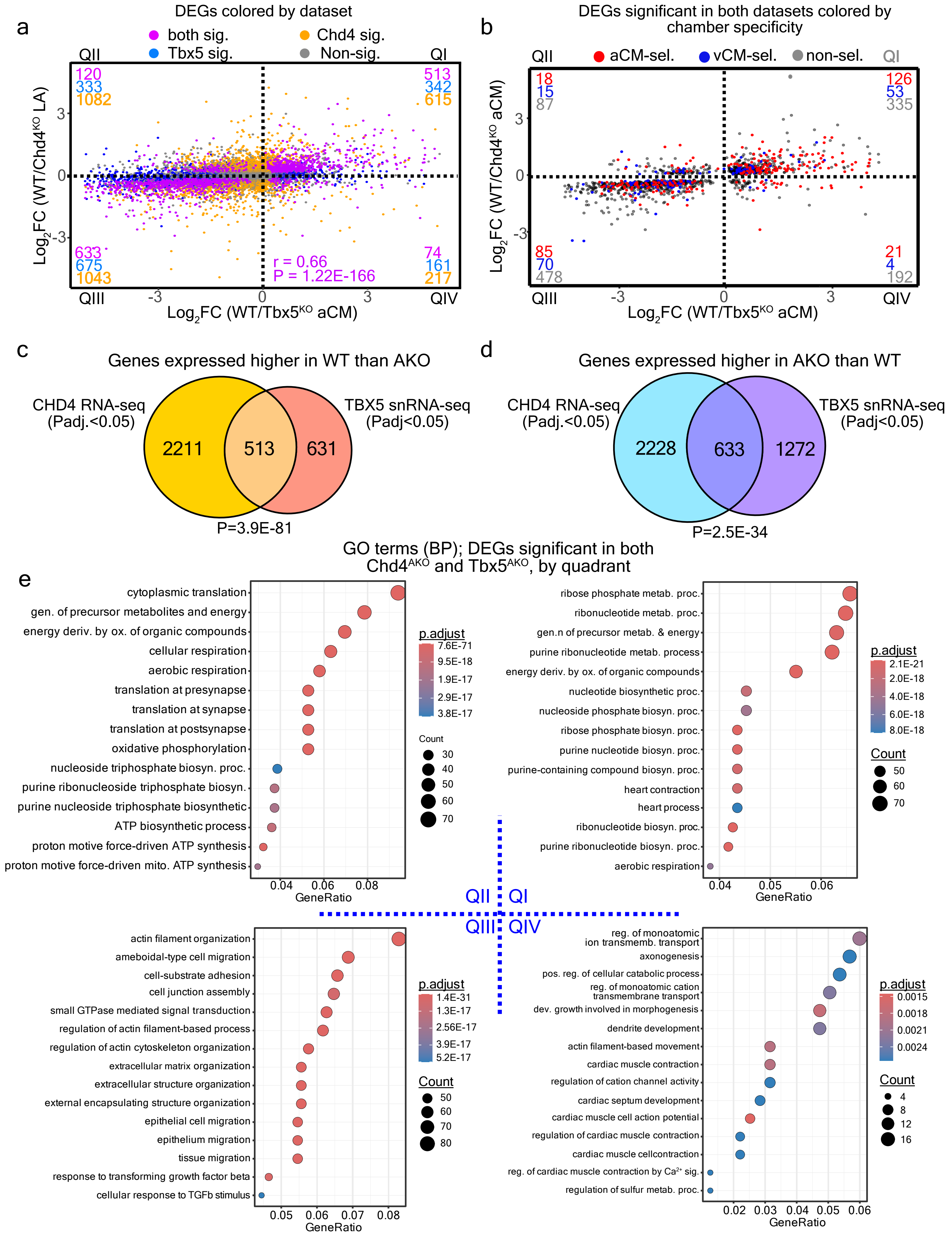


Supp. Fig. 2. CHD4 and TBX5 cooperate in the regulation of aCM genes

**Supp. Fig. 2. CHD4 and TBX5 cooperate in the regulation of aCM genes.**

**a,** Correlation between differential gene expression in $Tbx5^{AKO}$ and $Chd4^{AKO}$. Each gene was plotted by its fold-change between WT vs. $Tbx5^{AKO}$ and WT vs. $Chd4^{AKO}$. Genes were colored by the adjusted P value for each experiment. The Pearson correlation coefficient and p value are shown. **b,** Genes with significantly altered expression in both experiments were colored by aCM or vCM-selective expression. Most aCM-selective genes are located in QI, i.e. their expression was upregulated by both CHD4 and TBX5. c-d, Summary of gene changes from both experiments. **c,** Genes activated by CHD4 or TBX5. d, Genes repressed by CHD4 or TBX5. Hypergeometric test. e, GO term analysis of DEGs in each quadrant.
