## Supplemental Figure 3 for "CHD4 Interacts With TBX5 to Maintain the Gene Regulatory Network of Postnatal Atrial Cardiomyocytes"

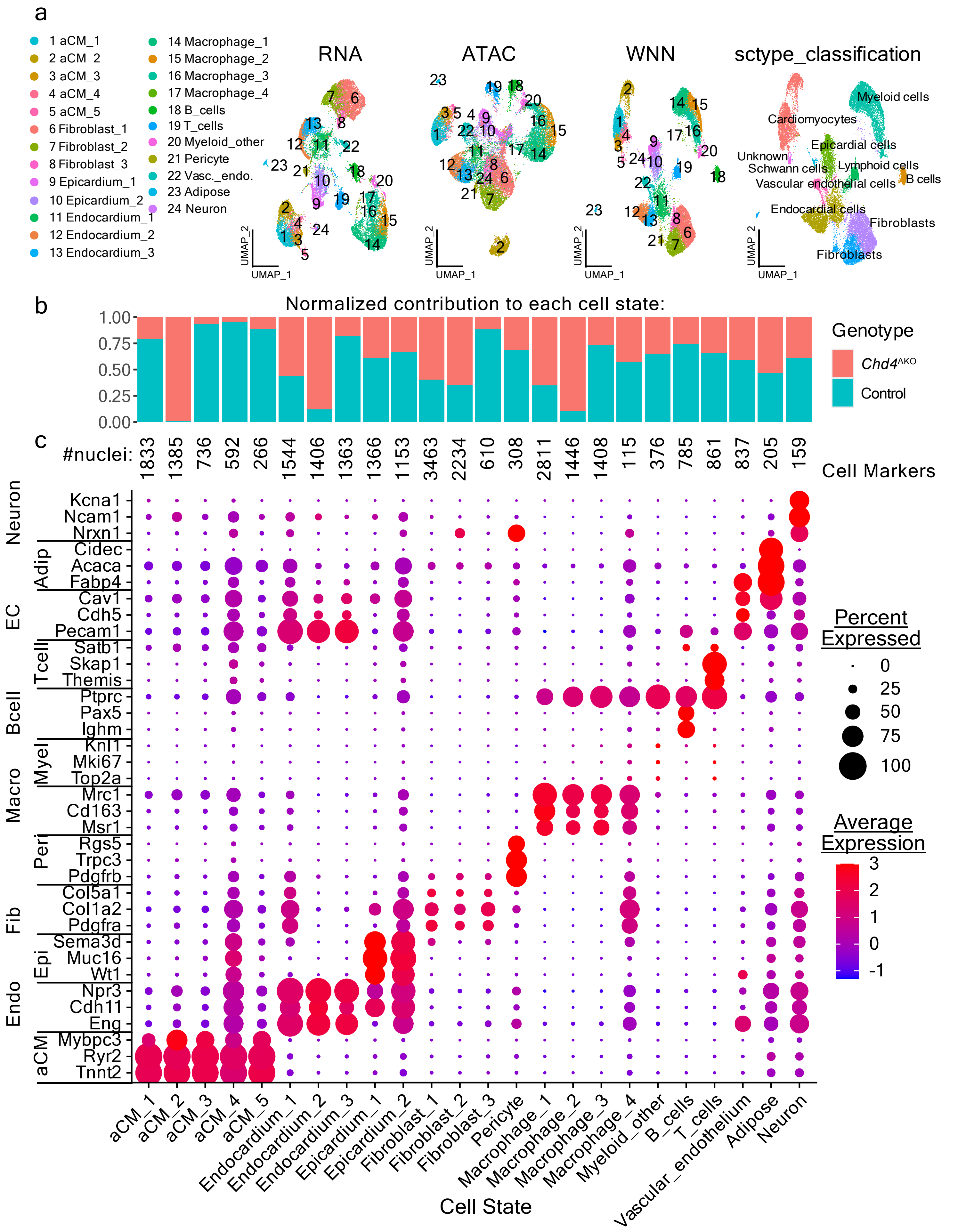


Supplemental Fig 3

Supp. Fig. 3. Concurrent snRNA-seq and snATAC-seq identifies distinct WT and Chd4$^{AKO}$ aCM clusters.

a, Cluster assignments and visualizations of the UMAP embeddings created for the RNA, ATAC and the WNN (weighted nearest neighbors) integration of the data types. Cell type classifications were assigyed using the SCtype package and the ‘Heart’ encyclopedia.b, Contribution of control and Chd4$^{AKO}$ samples to each cluster, normalized to the total number of cells from each group. c, Three curated marker genes for different atrial cell types. Numbers across the top of the plot indicate the number of nuclei of each type.
