## Supplemental Figure 4 for "CHD4 Interacts With TBX5 to Maintain the Gene Regulatory Network of Postnatal Atrial Cardiomyocytes"

**a**

### RNA data QC metrics

| Reaction | Sample | mean nFeature_RNA | median nFeature_RNA | mean nCount_RNA | median nCount_RNA | median fraction_mt |
|---|---|---|---|---|---|---|
| 1 | KO1 (KO) | 1645.42 | 1145 | 3859.56 | 1856 | 0.049 |
| 2 | KO2 (KO) | 1249.40 | 854 | 2477.31 | 1257 | 0.066 |
| 3 | S2 (KO) | 1366.30 | 991 | 2988.12 | 1577 | 0.074 |
| 4 | S4 (KO) | 1230.24 | 880 | 2491.55 | 1341 | 0.067 |
| 5 | WT1 (WT) | 1462.30 | 977 | 3330.38 | 1518 | 0.079 |
| 6 | WT2 (WT) | 1754.08 | 1061 | 4321.05 | 1680 | 0.083 |

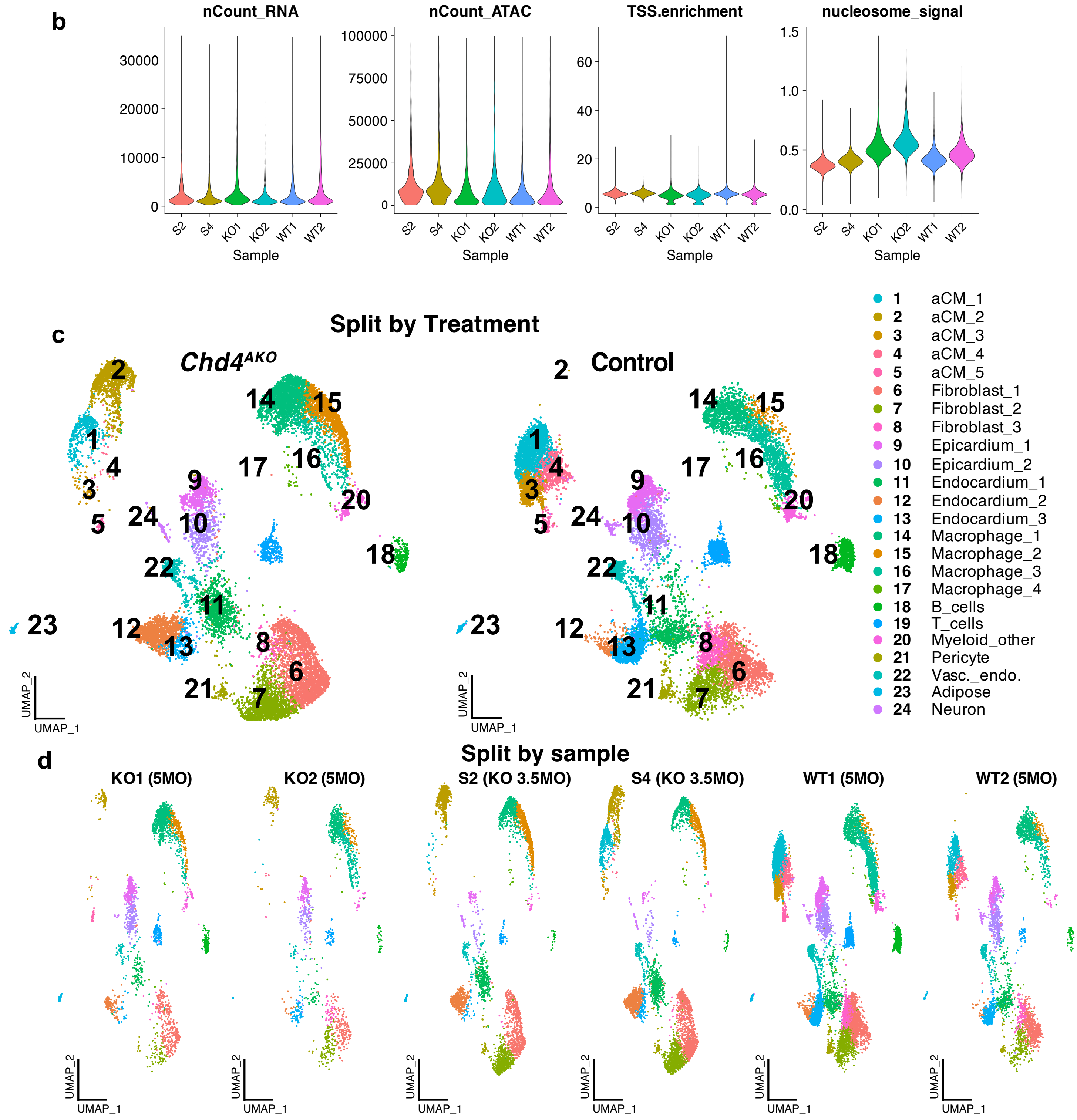


**Supplemental Fig 4. Quality control of the multiomics assay**

**Supp. Fig. 4. Quality control of the multiomics assay.**

**a**, Metrics for each individual 10x lane. These metrics were determined for the filtered snRNA-seq dataset. **b**, QC metrics for the filtered snATAC-seq dataset. **c**, The weighted nearest neighbor (WNN) UMAP embedding split by sample treatment. KO samples were labeled KO1, KO2, S2 and S4. WT samples were labeled WT1 and WT2. The UMAPs are colored by cluster identity. **d**, The WNN UMAP embedding split by sample. aCM_2 nuclei were almost exclusively derived from KO samples, which contain few aCM_1 nuclei.
