## Supplemental Figure 5 for "CHD4 Interacts With TBX5 to Maintain the Gene Regulatory Network of Postnatal Atrial Cardiomyocytes"

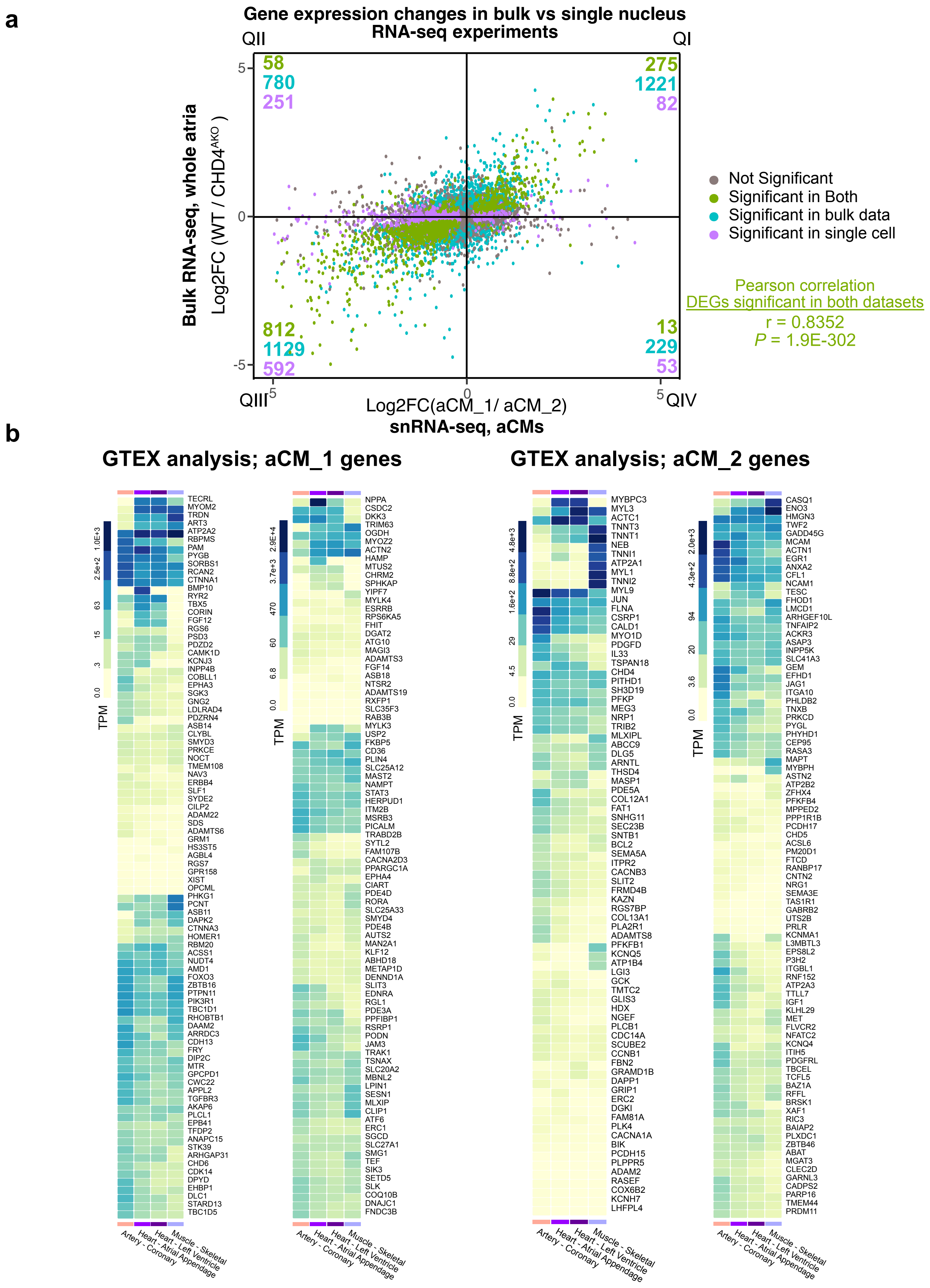


**Supplemental Fig 5.**

**Supp. Fig. 5. Examining DEGs of aCM_1 and aCM_2.**

**a**, Comparison of differential gene expression between control and *Chd4*$^{AKO}$ aCMs by snRNA-seq and control and *Chd4*$^{AKO}$ atria by bulk RNAseq. Points were colored by their significance in each assay. The Pearson correlation coefficient was calculated for significant DEGs from both assays (r=0.8352, P=1.9E-302). **b**, Top ranked DEGs from the control vs. *Chd4*$^{AKO}$ aCM snRNA-seq comparison were examined for expression levels in coronary arteries (enriched for smooth muscle cells), heart (atrial appendage), heart (left ventricle) and skeletal muscle, obtained from the GTEX RNAseq database. Genes more highly expressed in control than *Chd4*$^{AKO}$ aCMs tended to be highly expressed in heart tissue, while a subset of genes more highly expressed in *Chd4*$^{AKO}$ than control aCMs showed upregulation in skeletal muscle and coronary arteries.
