## Supplemental Figure 6 for "CHD4 Interacts With TBX5 to Maintain the Gene Regulatory Network of Postnatal Atrial Cardiomyocytes"

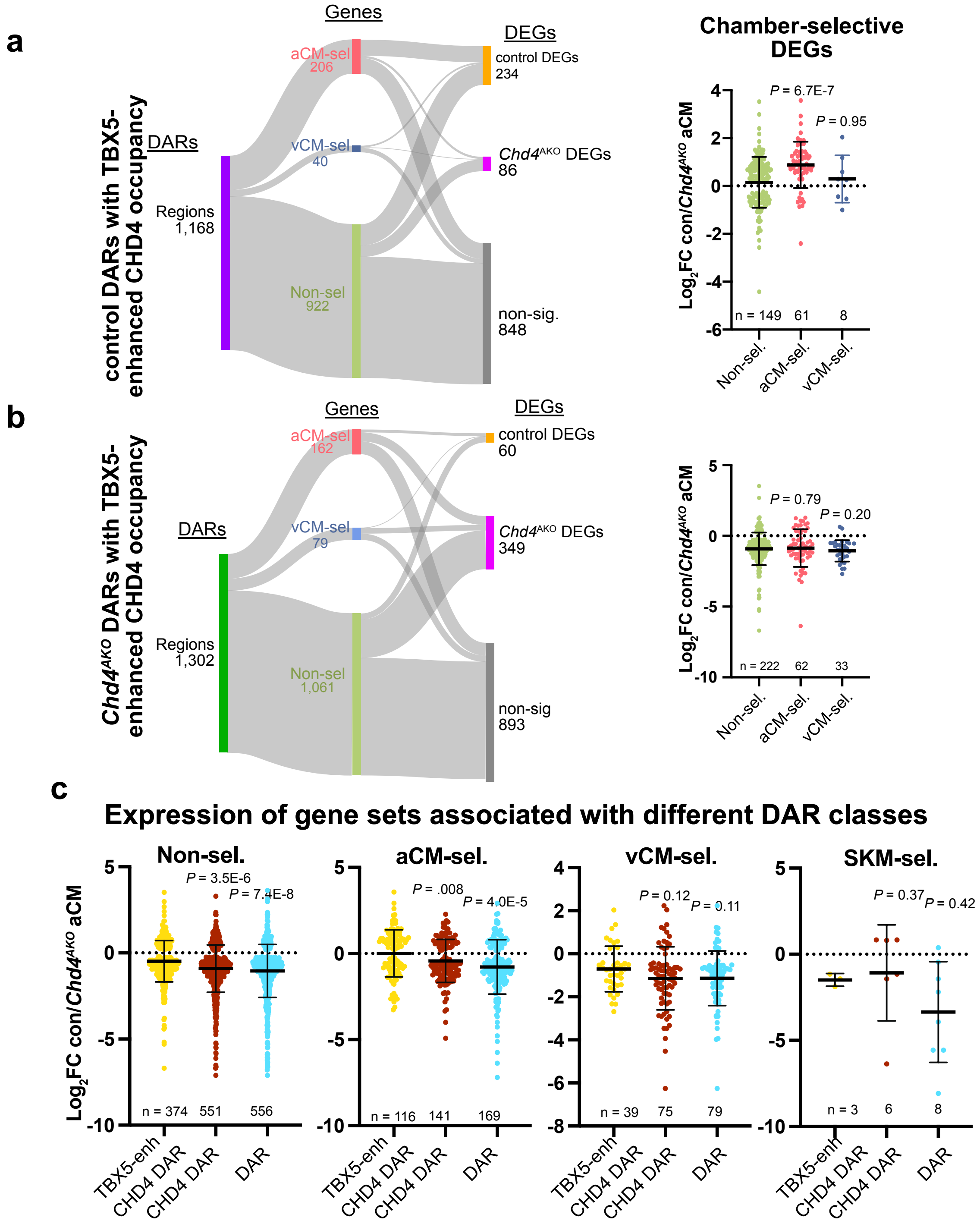


**Supplemental Fig 6. Gene targets of different DAR classes**

**Supplemental Fig 6. Gene targets of different DAR classes**

**a-b,**Control or *Chd4*$^{AKO}$ DARs with TBX5-enhanced CHD4 occupancy were classified as neighboring aCM-sel. genes, vCM-sel. genes, or non-sel. genes that did not show enrichment in either aCMs or vCMs. Regions were further classified by association with differentially expressed genes that were upregulated in control aCMs, *Chd4*$^{AKO}$ aCMs, or non-DEGs. The expression changes of Non-sel., aCM-sel., and vCM-sel. gene sets in control aCMs compared to *Chd4*$^{AKO}$ aCMs is shown on the righthand side of the figure panel. Wilcoxon test. The number of DEGs in each category is given along the X axis of the plot. Errors bars represent mean +/- SD.

**c,** Total expression changes of DEGs associated with TBX5-enhanced CHD4 bound DARs, CHD4 bound DARs, or DARs which were not bound by either protein. Wilcoxon test. The number of genes in each set is given along the X axis of the plot. Error bars represent mean+/- SD.
