## Supplemental Figure 7 for "CHD4 Interacts With TBX5 to Maintain the Gene Regulatory Network of Postnatal Atrial Cardiomyocytes"

**a**

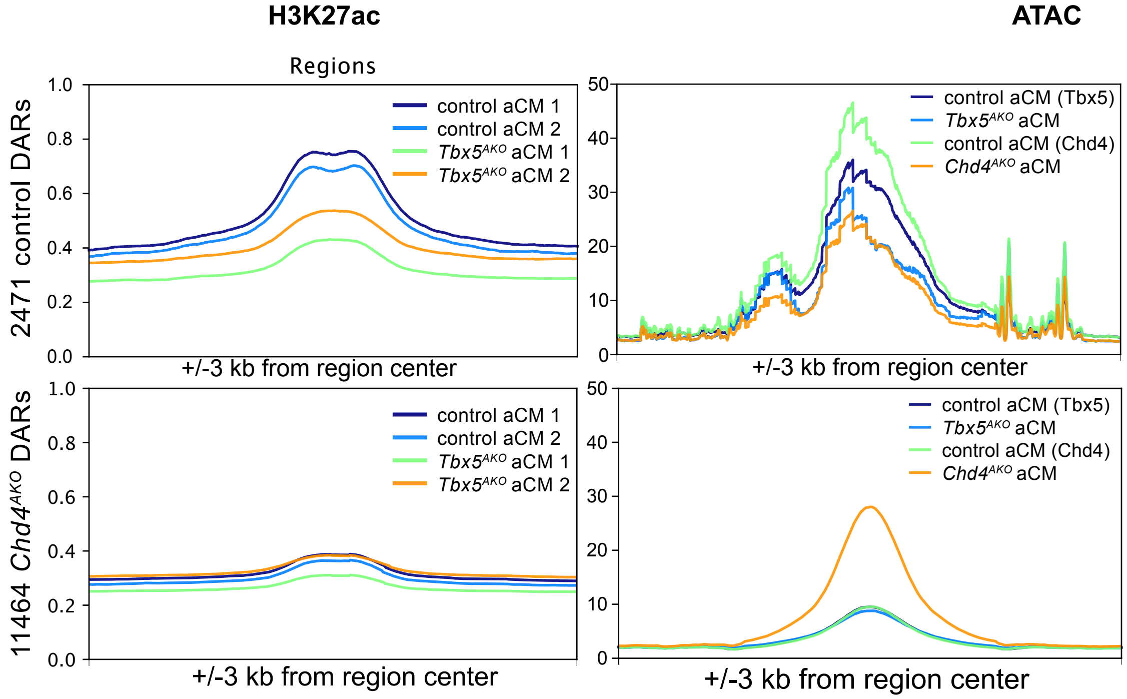


**b**

| Motif.Name | P.val(aCM1) | q.value.Benj._aCM1 | %targ._aCM1 | %backg._aCM1 | P.val(aCM2) | Q.value.Benj._aCM2 | %targ._aCM2 | %targ._aCM2 | Type |
|---|---|---|---|---|---|---|---|---|---|
| Mef2c(MADS)/GM12878-Mef2c-ChIP-Seq(GSE32465)/Homer | 1.00E-72 | 0 | 23.80% | 6.96% | 1.00E-40 | 0 | 15.98% | 5.61% | Shared |
| Mef2b(MADS)/HEK293-Mef2b.V5-ChIP-Seq(GSE67450)/Homer | 1.00E-67 | 0 | 34.85% | 14.40% | 1.00E-40 | 0 | 26.04% | 12.25% | Shared |
| Mef2d(MADS)/Retina-Mef2d-ChIP-Seq(GSE61391)/Homer | 1.00E-59 | 0 | 13.70% | 2.82% | 1.00E-41 | 0 | 9.83% | 2.30% | Shared |
| Mef2a(MADS)/HL1-Mef2a.biotin-ChIP-Seq(GSE21529)/Homer | 1.00E-55 | 0 | 20.98% | 6.80% | 1.00E-31 | 0 | 14.82% | 5.81% | Shared |
| NF1-halfsite(CTF)/LNCaP-NF1-ChIP-Seq(Unpublished)/Homer | 1.00E-38 | 0 | 61.47% | 42.48% | 1.00E-26 | 0 | 57.45% | 42.70% | Shared |
| Tgif1(Homeobox)/mES-Tgif1-ChIP-Seq(GSE55404)/Homer | 1.00E-25 | 0 | 77.83% | 63.41% | 1.00E-04 | 0 | 65.21% | 59.42% | Shared |
| Tgif2(Homeobox)/mES-Tgif2-ChIP-Seq(GSE55404)/Homer | 1.00E-25 | 0 | 80.22% | 66.30% | 1.00E-07 | 0 | 69.97% | 62.87% | Shared |
| NF1(CTF)/LNCAP-NF1-ChIP-Seq(Unpublished)/Homer | 1.00E-24 | 0 | 19.86% | 9.81% | 1.00E-57 | 0 | 26.96% | 10.87% | Shared |
| Meis1(Homeobox)/MastCells-Meis1-ChIP-Seq(GSE48085)/Homer | 1.00E-21 | 0 | 57.11% | 42.95% | 1.00E-05 | 0 | 47.85% | 41.43% | Shared |
| HIC1(Zf)/Treg-ZBTB29-ChIP-Seq(GSE99889)/Homer | 1.00E-19 | 0 | 62.84% | 49.55% | 1.00E-05 | 0 | 57.91% | 51.92% | Shared |
| GRE(NR),IR3/RAW264.7-GRE-ChIP-Seq(Unpublished)/Homer | 1.00E-36 | 0 | 16.27% | 5.82% | 1 | 0.2716 | 5.76% | 5.14% | Control_enriched |
| GRE(NR),IR3/A549-GR-ChIP-Seq(GSE32465)/Homer | 1.00E-30 | 0 | 10.96% | 3.32% | 1 | 0.186 | 3.69% | 3.06% | Control_enriched |
| PGR(NR)/EndoStromal-PGR-ChIP-Seq(GSE69539)/Homer | 1.00E-29 | 0 | 14.13% | 5.26% | 1 | 0.5428 | 4.45% | 4.29% | Control_enriched |
| ARE(NR)/LNCAP-AR-ChIP-Seq(GSE27824)/Homer | 1.00E-22 | 0 | 17.04% | 8.02% | 1 | 0.5608 | 7.30% | 7.12% | Control_enriched |
| PR(NR)/T47D-PR-ChIP-Seq(GSE31130)/Homer | 1.00E-15 | 0 | 65.75% | 53.90% | 1 | 0.7487 | 48.23% | 48.58% | Control_enriched |
| MRE(NR)/Neuro2A-NR3C2-ChIPnexus(GSE115417)/Homer | 1.00E-14 | 0 | 52.83% | 41.65% | 1 | 0.7775 | 38.79% | 39.23% | Control_enriched |
| ZNF91(Zf)/HEK-ZNF91.HA-ChIP-Seq(GSE162571)/Homer | 1.00E-06 | 0 | 25.34% | 19.37% | 1 | 0.4116 | 24.81% | 24.09% | Control_enriched |
| Smad4(MAD)/ESC-SMAD4-ChIP-Seq(GSE29422)/Homer | 1.00E-06 | 0 | 48.97% | 42.01% | 1 | 0.6175 | 41.71% | 41.57% | Control_enriched |
| SCL(bHLH)/HPC7-Scl-ChIP-Seq(GSE13511)/Homer | 1.00E-05 | 0 | 88.44% | 83.46% | 0.1 | 0.05 | 84.79% | 82.72% | Control_enriched |
| AMYB(HTH)/Testes-AMYB-ChIP-Seq(GSE44588)/Homer | 1.00E-05 | 0 | 41.52% | 34.92% | 0.1 | 0.0872 | 34.72% | 32.49% | Control_enriched |
| Fos(bZIP)/TSC-Fos-ChIP-Seq(GSE110950)/Homer | 0.1 | 0.0601 | 13.53% | 11.44% | 1.00E-37 | 0 | 22.58% | 10.27% | KO-enriched |
| Etv2(ETS)/ES-ER71-ChIP-Seq(GSE59402)/Homer | 0.1 | 0.0913 | 24.83% | 22.37% | 1.00E-10 | 0 | 30.88% | 22.88% | KO-enriched |
| NF-E2(bZIP)/K562-NFE2-ChIP-Seq(GSE31477)/Homer | 0.1 | 0.1072 | 1.97% | 1.28% | 1.00E-08 | 0 | 3.30% | 1.16% | KO-enriched |
| Atf3(bZIP)/GBM-ATF3-ChIP-Seq(GSE33912)/Homer | 0.1 | 0.1197 | 15.15% | 13.29% | 1.00E-41 | 0 | 25.50% | 11.80% | KO-enriched |
| Fra2(bZIP)/Striatum-Fra2-ChIP-Seq(GSE43429)/Homer | 0.1 | 0.1207 | 11.22% | 9.60% | 1.00E-39 | 0 | 20.81% | 8.80% | KO-enriched |
| Rfx5(HTH)/GM12878-Rfx5-ChIP-Seq(GSE31477)/Homer | 0.1 | 0.1246 | 10.79% | 9.21% | 1.00E-11 | 0 | 14.44% | 8.57% | KO-enriched |
| ERG(ETS)/VCaP-ERG-ChIP-Seq(GSE14097)/Homer | 0.1 | 0.134 | 39.73% | 37.21% | 1.00E-10 | 0 | 46.77% | 37.91% | KO-enriched |
| AP-1(bZIP)/ThioMac-PU.1-ChIP-Seq(GSE21512)/Homer | 0.1 | 0.1577 | 17.38% | 15.56% | 1.00E-40 | 0 | 28.19% | 13.94% | KO-enriched |
| JunB(bZIP)/DendriticCells-Junb-ChIP-Seq(GSE36099)/Homer | 0.1 | 0.159 | 12.67% | 11.09% | 1.00E-40 | 0 | 22.73% | 9.99% | KO-enriched |
| Nur77(NR)/K562-NR4A1-ChIP-Seq(GSE31363)/Homer | 0.1 | 0.1724 | 4.54% | 3.60% | 1.00E-04 | 0 | 5.45% | 3.16% | KO-enriched |

**Supplemental Fig 7. H3K27ac levels and motif enrichment of control and KO regions**

**Supp. Fig. 7. Characterization of *Chd4*$^{AKO}$ and control DARs. a,** We analyzed H3K27ac occupancy of *Chd4*$^{AKO}$ and control DARs using our previous H3K27ac HiChIP data from *Tbx5*$^{AKO}$ and control atria processed as regular ChIP-seq (GSE222970). H3K27ac was depleted at control DARs in *Tbx5*$^{AKO}$ aCMs. **b**, Motifs over-represented in DARs. All of the otifs selectively enriched in control DARs and the top 10 motifs selectively enriched in *Chd4*$^{AKO}$ DARs are shown.
